# A two-state photoconversion model predicts the spectral response dynamics of optogenetic systems

**DOI:** 10.1101/081430

**Authors:** Evan J. Olson, Constantine N. Tzouanas, Jeffrey J. Tabor

## Abstract

In optogenetics, light signals are used to control genetically engineered photoreceptors, and in turn manipulate biological pathways with unmatched precision. Recently, evolved photoreceptors with diverse *in vitro*-measured wavelength and intensity-dependent photoswitching properties have been repurposed for synthetic control of gene expression, proteolysis, and numerous other cellular processes. However, the relationship between the input light spectrum and *in vivo* photoreceptor response dynamics is poorly understood, restricting the utility of these optogenetic tools. Here, we advance a classic *in vitro* two-state photoreceptor model to reflect the *in vivo* environment, and combine it with simplified mathematical descriptions of signal transduction and output gene expression through our previously engineered green/red and red/far red photoreversible bacterial two-component systems (TCSs). Additionally, we leverage our recent open-source optical instrument to develop a workflow of spectral and dynamical characterization experiments to parameterize the model for both TCSs. To validate our approach, we challenge the model to predict experimental responses to a series of complex light signals very different from those used during parameterization. We find that the model generalizes remarkably well, predicting the results of all categories of experiments with high quantitative accuracy for both systems. Finally, we exploit this predictive power to program two simultaneous and independent dynamical gene expression signals in bacteria expressing both TCSs. This multiplexed gene expression programming approach will enable entirely new studies of how metabolic, signaling, and decision-making pathways integrate multiple gene expression signals. Additionally, our approach should be compatible with a wide range of optogenetic tools and model organisms.

**Significance statement:** Light-switchable signaling pathways (optogenetic tools) enable precision studies of how biochemical networks underlie cellular behaviors. We have developed a versatile mathematical model based on a two-state photoconversion mechanism that we have applied to the *E. coli* CcaSR and Cph8-OmpR optogenetic tools. This model enables accurate prediction of the gene expression response to virtually any light source or mixture of light sources. We express both optogenetic tools in the same cell and apply our model to program two simultaneous and independent gene expression signals in the same cell. This method can be used to study how biological pathways integrate multiple inputs and should be extensible to other optogenetic tools and host organisms.

## Introduction

Most optogenetic tools are based on a photoreceptor protein with a light-sensing domain that regulates an effector domain, which in turn generates a biological signal such as gene expression. One can consider a simplified model wherein a photoreceptor is produced in a ‘ground’ state and switched to an ‘active’ state by activating wavelengths (i.e. forward photoconversion)^1^. Active state photoreceptors thermally revert to the ground state with a characteristic timescale that ranges from milliseconds^2^ to more than a month^3^. Certain photoreceptors, exemplified by the linear tetrapyrrole (bilin)-binding phytochrome (Phy) and cyanobacteriochrome (CBCR) families are also photoreversible where reversion from the active to ground state is driven by deactivating wavelengths^4–6^.

Two-component systems (TCSs) are signal transduction pathways that control gene expression and other processes in response to chemical or physical stimuli (inputs). Canonical TCSs comprise two proteins; a sensor histidine kinase (SK) and a response regulator (RR). The SK is produced in a ground state, which often has low kinase activity toward the RR. When it detects an input via a N-terminal sensing domain, the SK uses ATP to autophosphorylate on a histidine residue within a C-terminal kinase domain. This phosphoryl group is then transferred to an aspartate on the RR. In most cases the phosphorylated RR (RR~P) binds to a target promoter, activating transcription. Many SKs are bi-functional and the kinase domain dephosphorylates the RR~P in the absence of the input or presence of a different, de-activating input.

We have previously engineered two spectrally distinct photoreversible *E. coli* TCSs, CcaSR and Cph8-OmpR^7–9^. CcaS is a SK with a CBCR sensing domain that absorbs light via a covalently ligated phycocyanobilin (PCB) chromophore produced by an engineered metabolic pathway. Holo-CcaS is produced in an inactive, green light sensitive ground state, termed Pg, with low kinase activity. Upon green light exposure, CcaS Pg switches to a red light sensitive active state (Pr) with high kinase activity toward the RR CcaR. CcaR~P binds to the promoter P_*cpcG2*−172_, activating transcription. Red light drives CcaS Pr to revert to Pg. Cph8 is a chimeric SK containing the PCB-binding Phy light-sensing domain of *Synechocystis* PCC6803 Cph1 and the signaling domain of *E. coli* EnvZ. In contrast to CcaS, Cph8 has high kinase activity toward the *E. coli* RR OmpR in the ground state (Pr) and low kinase (high phosphatase) activity in a far-red absorbing activated state (Pfr). OmpR~P binds and activates transcription from the P_*ompF*146_ promoter. Data from our group and others suggest that CcaS Pr is stable for hours or more^10,11^ while Cph8 Pfr is far less stable^11^.

Recently, we developed predictive phenomenological models of the responses of CcaSR and Cph8-OmpR to green and red light intensity signals, respectively^11^. These models describe a three step dynamical response comprising a pure delay, an intensity-dependent first-order transition in output gene expression rate, and a first-order transition in the concentration of the output gene set by cell growth rate. By measuring the expression of a reporter gene over time in response to a series of light step-changes of different initial and final intensities, we parameterized these three timescales for both light sensors.

Next, we used these models to program tailor-made gene expression signals with an unrivaled degree of control and predictability^11^. In particular, we combined the models with a custom ‘light program generator’ algorithm that accepts a reference (desired) gene expression signal as an input and produces a green or red light signal that drives CcaSR or Cph8-OmpR to produce that gene expression output experimentally. We utilized this ‘biological function generator’ method to create linear ramps and sine waves of a transcriptional repressor in order to characterize the input/output dynamics of a synthetic gene circuit^11^.

Despite their utility, our previous models have several key limitations. First, they can only predict the responses of the optogenetic tools to the specific light sources used during parameterization. Second, they cannot account for perturbations introduced by secondary light sources such as those that might be used for simultaneous measurement of fluorescent reporter proteins or multiplexed control of both tools in the same cell. Third, the models yield few insights into the mechanistic origin of the observed response dynamics. For example, the models captured, but could not elucidate the origin of, our observation that the rate of the gene expression transition depends upon the direction and final intensity of the light step change.

An *in vitro* two-state model^1,12,13^ describing the intensity and wavelength dependence of switching between ground and active states has previously been used to describe photoswitching of Phys^1^, CBCRs^3^, bacteriophytochromes^13^, LOV domains^14^, and Cryptochromes^15^ among others. In this model, the sensors are characterized by their ground- and active-state photoconversion cross sections (PCSs), *σ*_g_(*λ*) and *σ*_a_(*λ*), which enable direct calculation of the forward and reverse photoconversion rates, *k*_1_ and *k*_2_, in response to photons of wavelength *λ*. The PCS, expressed as the photoconversion rate per unit light intensity (min^−1^ [µmol m^−2^ s^−1^]^−1^), is proportional to the product of *ε*(*λ*), the molar extinction coefficient (m^2^ mol^−1^), which is related to the probability of the photoreceptor absorbing a photon, and the *ϕ*(*λ*), the quantum yield (unitless), which describes the probability of photoconversion upon photon absorption. Given knowledge of both PCSs (*σ*_i_(*λ*)), one can compute both photoconversion rates (*k*_i_) for a light source with a known spectral flux density *n*_light_(*λ*) (µmol m^−2^ s^−1^ nm^−1^) by calculating the spectral overlap integral *k*_*i*_ = *∫ σ*_*i*_ · *n*_light_ *dλ*. The photoconversion rates can then be used to calculate the populations of ground and active state photoreceptor.

Despite its potential for predicting photoreceptor responses to virtually any light condition, the two-state model has not been explored for optogenetics. In particular, the complete *σ*_i_(*λ*) has not been determined for any photoreceptor used in optogenetics. Even if *σ*_i_(*λ*) were to be determined, the two-state model would need to be extended to capture photoreceptor production and decay dynamics in the *in vivo* environment. Finally, an additional model would be needed to capture the biological events that occur downstream of the photoreceptor.

Here, we extend the two-state model for the *in vivo* environment, develop a new strategy for estimating *σ*_i_(*λ*) *in vivo,* and combine these efforts with simplified descriptions of TCS signaling and gene expression for CcaSR and Cph8-OmpR. We then develop a standard set of spectral and dynamic characterization experiments to parameterize the overall TCS photoswitching model. We validate the models by predicting and measuring the gene expression response of both systems to spectrally and dynamically diverse light programs. Finally, we express CcaSR and Cph8-OmpR in the same cell and combine the models with our biological function generator approach to achieve the first multiplexed programming of gene expression dynamics.

## Results

### TCS photoconversion model

We constructed a TCS ‘sensing model’ (**Methods**) by adding terms for production of new ground state photoreceptors (*S*_*g*_) at rate *k*_*S*_ and dilution of *S*_g_ and active state photoreceptors (*S*_*a*_) at rate *k*_dil_ to the two-state model (**Fig. 1a**). The sensing model accepts any *n*_light_(*λ*) input and produces *S*_g_ and *S*_a_ populations as an output (**Fig. 1b,c**). The ratio *S*_a_/*S*_g_ feeds into an ‘output model’ comprising a phenomenological description of TCS signaling and a standard model of output gene expression (**Fig. 1c**). The TCS signaling model (**Methods**) describes a pure time delay (*τ*) and Hill-function mapping 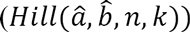 between *S*_a_/*S*_g_ and output gene production rate (*k*_*G*_). In our initial experiments, we utilize superfolder GFP (*G*) as the output and quantify its expression level in Molecules of Equivalent Fluorescein (MEFL). 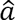 is the range of possible *k*_*G*_ values, 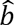 is the minimum value of *k*_*G*_, *n* is the Hill parameter, and *k* is *S*_a_/*S*_g_ ratio resulting in 50% maximal system response. Together, these terms capture SK autophosphorylation, phosphotransfer, RR dimerization, DNA binding, promoter activation, and *G* production. *G* is degraded in a first-order process with rate *k*_dil_ (**Methods**), and has a minimum concentration 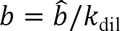 and concentration range 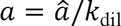 given a constant cell growth rate.

**Fig 1.**
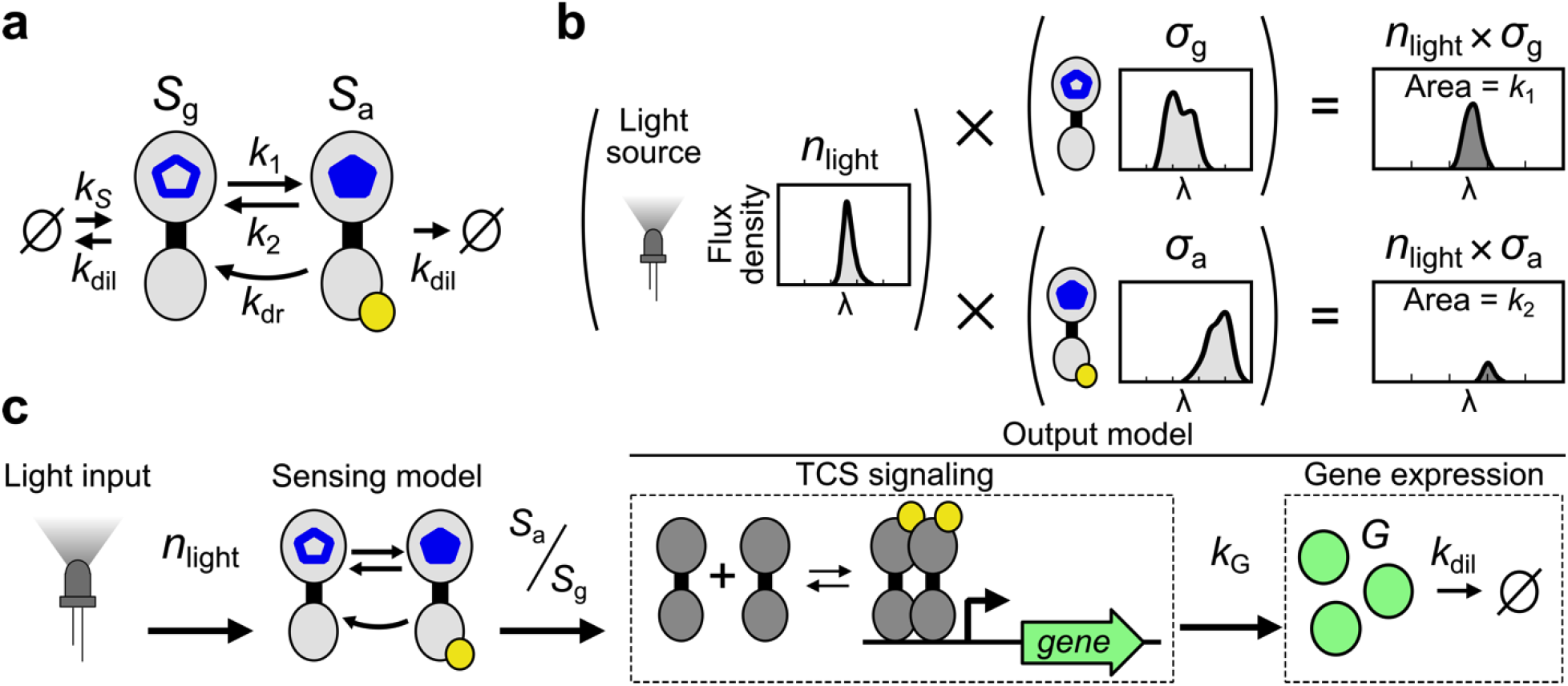
TCS photoconversion model. (a) The two-state photoreceptor model, which includes ground (*S*_g_) and active (*S*_a_) state photoreceptors (aka sensors), photoconversion rates *k*_1_ and *k*_2_, and dark reversion rate *k*_dr_, is converted to a “sensing model” for *in vivo* environments by adding a *S*_g_ production rate *k*_*S*_ that captures both gene expression and holo-protein formation, and a dilution rate *k*_dil_ due to cell growth and sensor degradation (**Methods**). The hollow blue pentagon represents a chromophore in the ground state while the filled blue pentagon represents that in the activated state. (b) Photoconversion rates are determined by the overlap integral of the spectral flux density of the light source (*n*_light_) and the *S*_g_ and *S*_a_ photoconversion cross sections *σ*_g_ and *σ*_a_ (**Methods**). (c) The sensing model converts *n*_light_ into the active ratio of light sensors S_a_/S_g_ which feeds into an “output model” with a simplified model of TCS signaling that regulates the production rate *k*_*G*_ of the target protein *G*, which is diluted due to cell growth and proteolysis as *k*_dil_ (**Methods**).

### Light source model

Most light sources have a fixed spectral flux density (i.e. output spectrum) that scales with light intensity (*I*, µmol m^−2^ s^−1^). For such light sources, we can write 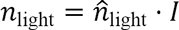 where 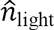 is the output spectrum at 1 µmol m^−2^ s^−1^. To quantify the overlap between *n*_light_ and *σ*_*i*_ for a given photoreceptor, we introduce 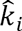 as the photoconversion rate per unit light intensity (min^−1^ [µmol m^−2^ s^−1^]^−1^). Then, for a given light source, 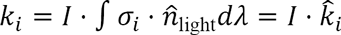. That is, *k*_1_ and *k*_2_ take on values proportional to light intensity.

### Dynamical and spectral characterization of CcaSR

We designed a set of four gene expression characterization experiments (**File S1-2, Note S1-2**) to train the TCS photoconversion model for CcaSR (**Fig. 2a**). First, we quantify activation dynamics by preconditioning *E. coli* expressing CcaSR (**Fig. S1**) in the dark, introducing step increases in green light (centroid wavelength *λ*_*c*_ = 526 nm, **Table S1–3, File S3-4, Note S3**) to different intensities, and measuring sfGFP levels over time by flow cytometry (**Methods, Fig. 2b, S2**). Second, we measure de-activation dynamics by preconditioning the cells in different intensities of green light and measuring the response to step decreases to dark (**Fig. 2c, S2**). Third, we measure the ground state spectral response by exposing the bacteria to 23 LEDs with *λ*_*c*_ spanning 369 to 958 nm at over three orders of magnitude intensity (**Methods, Fig. 2d, S3, Table S1–3, File S3-4, Note S3**) and measuring sfGFP at steady state. Finally, we measure the activated state spectral response by repeating the previous experiment in the presence of a constant intensity of activating light (**Fig. 2e, S3**).

**Fig 2.**
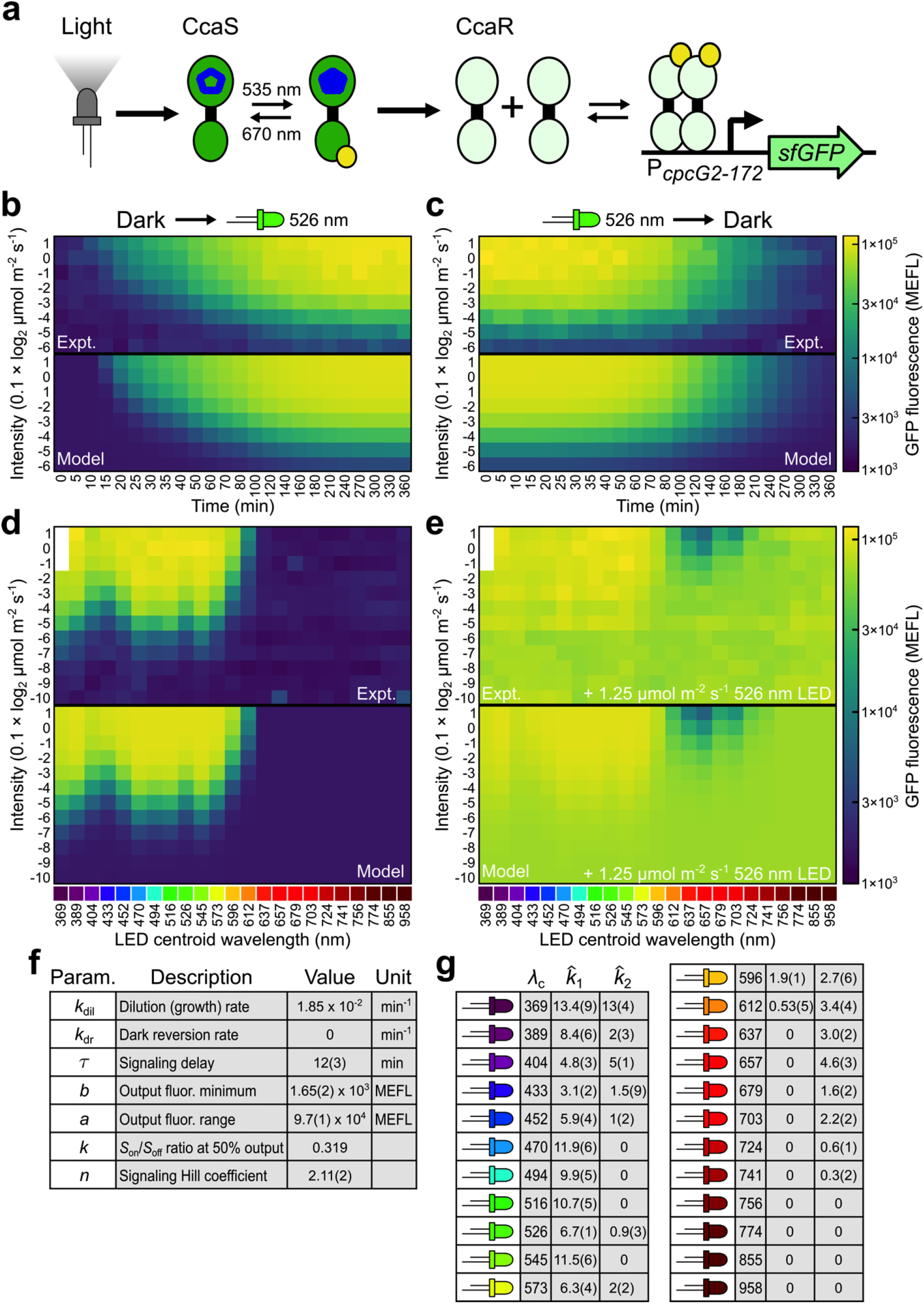
Characterization and model parameterization for CcaSR. (a) Schematic of CcaSR TCS with sfGFP output. Wavelength values represent *in vitro* measured absorbance maxima. (b-e) Training data for the full CcaSR system model (**Fig. 1c**). Experimental observations (“Expt.”) and simulations of the best-fit model (“Model”) are shown for each set. In particular, the response dynamics to step (b) increases from dark to eight different intensities and (c) decreases from eight different intensities to dark were evaluated using the *λ*_c_ = 526 nm LED. Time points are distributed unevenly to increase resolution of the initial response. (c-d) Steady-state intensity dose-response to a set of 23 “spectral LEDs” with *λ*_c_ spanning 369 nm to 958 nm. (c) Forward photoconversion is primarily determined by the response to the spectral LEDs. (d) Reverse photoconversion is analyzed by including light from a second, activating LED (*λ*_c_ = 526 nm at 1.25 µmol m^−2^ s^−1^). The *λ*_c_ = 369 nm LED is not capable of reaching the brightest intensities, and thus those data points are not included. Light intensities are shown in units of 0.1 × log_2_ µmol m^−2^ s^−1^ scale (e.g. a value of 1 corresponds to 10 × 2^1^ = 20 µmol m^−2^ s^−1^). sfGFP fluorescence is calibrated to MEFL units (**Methods**). Each row of measurements in panels b-e was collected in a single 24-well plate. The 40 plates required to produce the training dataset were randomly distributed across eight LPAs over five separate trials (**Methods, File S1-2**). Each color patch represents the arithmetic mean of a single population of cells. (f-g) Best-fit model parameters produced via nonlinear regression of the model to training data (**Methods, Table S4**). 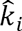 are unit photoconversion rates (i.e. 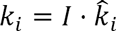, where *I* is the LED intensity in µmol m^−2^ s^−1^). Uncertainty in the least-significant digits are indicated in parenthesis.

### CcaSR model parameterization

We used nonlinear regression (**Methods, Table S4**) to fit the model to these data. While the resulting parameters recapitulate the known properties of the system (**Fig. 2f-g, S4**), the value of the Hill parameter *k* is weakly determined (**Table S4**). In particular, alterations in *k* from the best-fit value can be compensated for by changes in 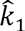 and 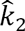 (**Fig. S5**). Thus, we cannot confidently determine the absolute rates of forward and reverse photoconversion. Nonetheless, fixing *k* at its best fit value results in model predictions that quantitatively agree with the experimental measurements (**Fig. 2b-e**). However, the ultimate validation of this approach involves predicting the response of CcaSR to a wide range of spectral and dynamical light inputs different from those used in parameterization.

### Spectral validation of the CcaSR photoconversion model

To predict the response of an optogenetic tool to a given light source, knowledge of *σ*_*i*_ is required. To estimate *σ*_*i*_ for CcaSR, we used non-linear regression to fit a cubic spline to the previously determined photoconversion rates for each of the 23 LEDs (**Methods Fig. 3a, Fig. S6–7**). Importantly, our regression procedure considers the response of CcaSR to the full spectral output of each LED, not just its centroid wavelength. To validate the resulting *σ*_*i*_ estimate, we measured 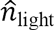 for a previously untested set of eight color-filtered white light LEDs designed to have complex spectral characteristics (**Table S1–3, File S3-4, Note S3**) and calculated an expected 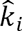 for each (**Fig. 3b**). In combination with the remaining model parameters (**Fig. 2f**), we used these 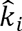 to predict the steady state intensity dose-response to these eight LEDs in the presence and absence of activating light (*λ*_*c*_ = 526 nm). These predictions are remarkably accurate for LEDs 1-5 (root-mean-square errors (RMSEs) from 0.11 to 0.18, **Methods**), which drive sfGFP to high levels, and 7 and 8, which drive low expression (RMSE = 0.14 and 0.18, respectively), but slightly less so for LED 6 (RMSE = 0.26), which drives sfGFP to an intermediate expression level (**Fig. 3c**).

**Fig 3.**
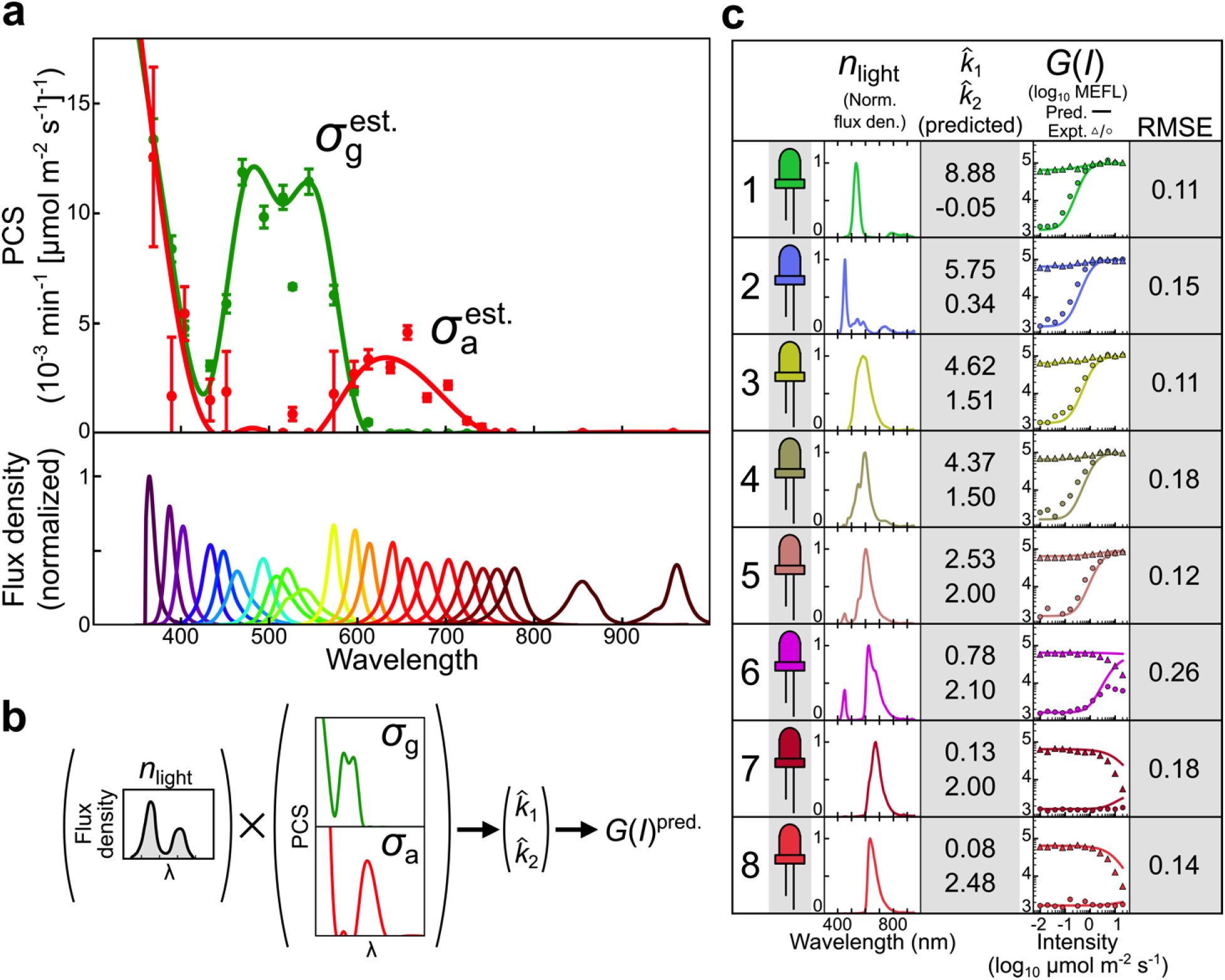
Estimation of the CcaS photoconversion cross section and spectral validation of the CcaSR model. (a) We estimate the continuous ground and active state PCSs of CcaS (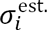, lines) by regressing cubic splines to minimize the difference between experimentally-determined photoconversion rates (points) and those calculated via 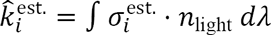 (**Methods, Fig. S6,7**). Error bars correspond to the standard error of the fits for the experimentally-determined photoconversion rates. The normalized spectral flux densities of the spectral LEDs are shown at bottom. (b) Using 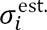 to predict photoconversion rates for light sources not in the spectral LED training set. Predicted photoconversion rates are integrated into the CcaSR model by keeping all other parameters (**Fig. 2f**) fixed, enabling prediction of the intensity dose-response of CcaSR to the new light source (i.e. *G*(*I*)^*pred.*^). (c) Spectral validation of the CcaSR model and 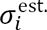 consists of prediction of the intensity dose-response for eight challenging, broad-spectrum light sources constructed by applying colored filters over white-light LEDs (**Methods, Table S1–3, File S1-4**). Measured *n*_light_, predicted 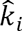, measured and predicted intensity dose-response curves, and RMSE between model and prediction are shown for each LED (**Methods**). The forward and reverse intensity responses are determined using the filtered LED alone (circles) and in the presence of a second activating LED (*λ*_*c*_ = 526 nm at 1.25 µmol m^−2^ s^−1^, triangles). The simulated responses are determined using the calculated photoconversion rates (**Methods**). RMSE relative errors are expressed in log_10_ decades (**Methods**). Data was collected across four LPAs, and the forward (circles) and reverse (triangles) intensity responses were collected over two separate experimental trials (**Methods, File S1-2**). Each data point represents the arithmetic mean of a single population of cells.

### Dynamic validation of the CcaSR photoconversion model

Our biological function generator method constitutes a rigorous validation of the predictive power of a model because the light inputs and gene expression outputs are temporally complex and cover a wide range of levels. To validate our CcaSR photoconversion model, we first designed a challenging reference gene expression signal (**Fig. 4, File S5**). The signal starts at *b* and then increases linearly (on a logarithmic scale) over 90% of the total CcaSR response range over 210 min. After a 60 min. hold, the signal decreases linearly to an intermediate expression level over another 210 min. Using this reference, we then used the model to computationally design four light time courses each with different LEDs or LED mixtures (**Methods, File S6**). “UV mono” utilizes a single UV LED (*λ*_*c*_ = 389 nm) (**Fig. 4a**) to demonstrate control of CcaSR with an atypical light source. “Green mono” uses the *λ*_*c*_ = 526 nm LED (**Fig. 4b**) to demonstrate predictive control with a typical light source. “Red perturbation” combines “Green mono” with a strong red (*λ*_*c*_ = 657 nm) sinusoidal signal (**Fig. 4c**) designed to demonstrate the perturbative effects of alternative light sources. Finally, in “Red compensation”, the “Green mono” time course is re-optimized to compensate for the impact of “Red perturbation” (**Fig. 4d, Methods**).

**Fig 4.**
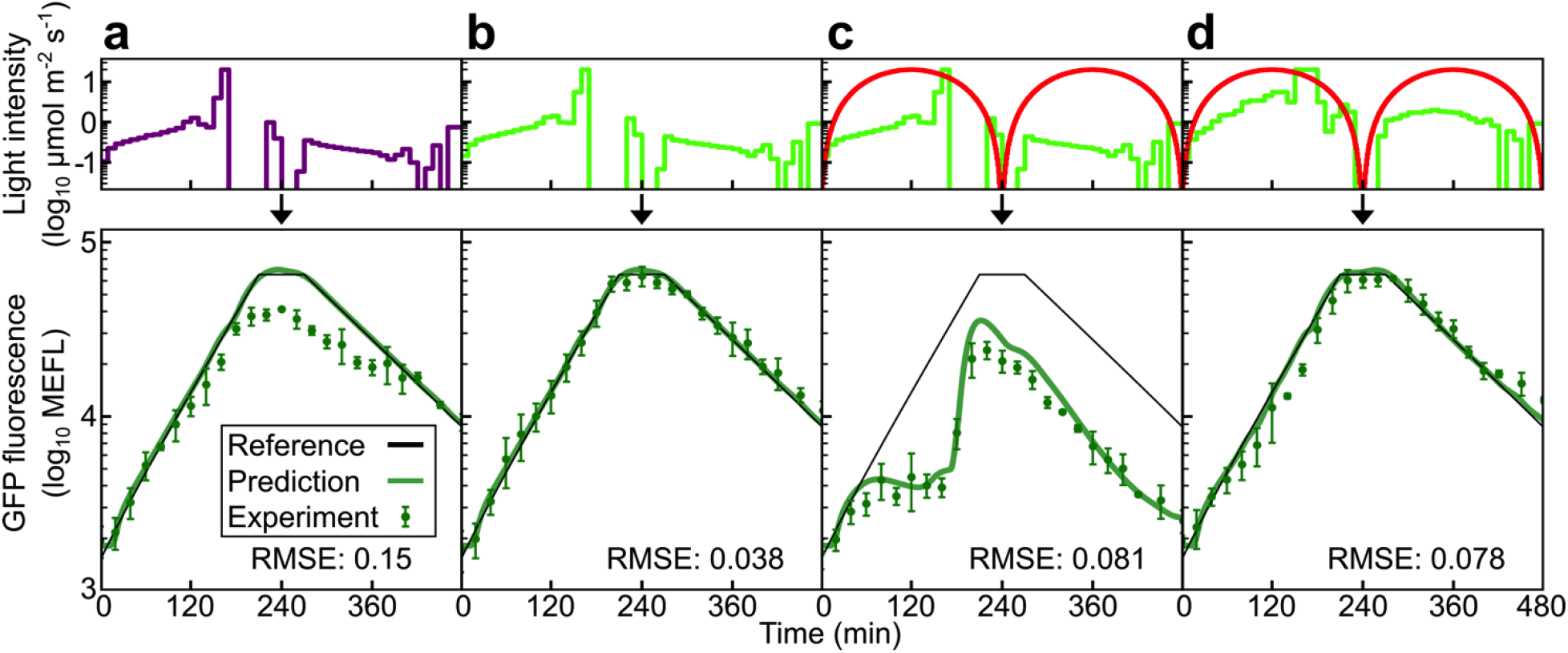
Dynamical validation of the CcaSR model. We compare model predictions of dynamical CcaSR sfGFP output to experimental measurements for time-varying light inputs from UV (*λ*_*c*_ = 389 nm), green (*λ*_*c*_ = 526 nm), or green plus red (*λ*_*c*_ = 657 nm) light. In all cases, the light program generator algorithm (LPG, **Methods**) is used to design light signals predicted to drive sfGFP to follow the reference *G*(*t*)_Ref._, consisting of a ramp up, hold, and ramp down on a logarithmic scale (**File S5-6**). (a) “UV mono”. The LPG-generated UV light signal drives the CcaSR system along a trajectory predicted to follow the reference signal. (b) “Green mono”. The green LED alone provides an optimized input signal. (c) “Red perturbation”. The green LED provides the “Green mono” signal, while the red LED generates a sinusoidal perturbative signal (center) with a 240-minute period and 20 µmol m^−2^ s^−1^ peak-to-peak amplitude. (d) “Red compensation”. The red perturbative signal is again present. However, the LPG redesigns the green light signal to account for its presence. Light signals are shown in units of log_10_ µmol m^−2^ s^−1^, and RMSE relative errors are expressed in log_10_ decades (**Methods**). Error bars correspond to the standard deviation in fluorescence measurements over three independent experimental trials (**Table S4**).

**Fig 5.**
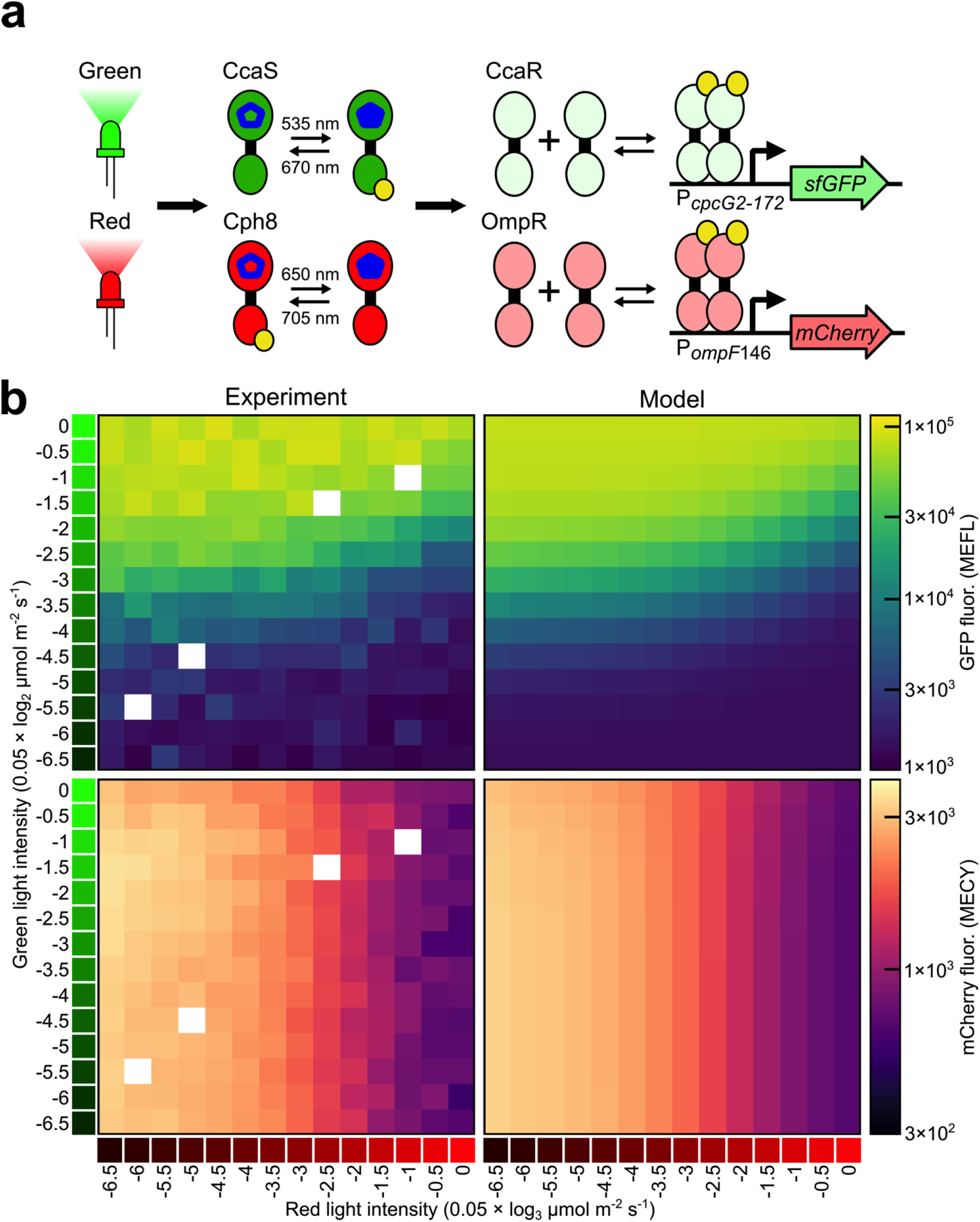
Characterization and modeling of a multiplexed CcaSR/Cph8-OmpR system. (a) CcaSR and Cph8-OmpR are co-expressed in a single strain. CcaSR regulates the expression of sfGFP, while Cph8-OmpR regulates the expression of mCherry. Wavelength values are as in **Fig. 2a** (b) Training data for the multiplexed model (“Experiment”, **File S8**) consists of a two-dimensional steady-state intensity dose-response to green (*λ*_*c*_ = 526 nm) and red (*λ*_*c*_ = 657 nm) light. The light intensities are logarithmically distributed, with the green light varying on a 0.05 × log_2_ µmol m^−2^ s^−1^ scale (e.g. a value of −1 corresponds to 20 × 2^−1^ = 10 µmol m^−2^ s^−1^) and the red light varying over a 0.05 × log_3_ µmol m^−2^ s^−1^ scale (e.g. a value of −1 corresponds to 20 × 3^−1^ = 6.67 µmol m^−2^ s^−1^). The different intensity ranges are used to maintain a high-resolution measurement despite the differences in the intensity dose-responses of the two systems. The four missing intensity values (white boxes) were not collected. The training data was used to re-fit the *a*, *b*, *n*, and *k* Hill function parameters for the CcaSR and Cph8-OmpR models (**Table S6**). Simulated steady-state responses to the same light environments for the best-fit dual-system models (**Table S6**) are shown (“Model”). mCherry fluorescence is calibrated to MECY units (Molecules of Equivalent Cy5, **Methods**). RMSE relative errors are expressed in log_10_ decades (**Methods**). Data was collected in one experimental trial, and the 192 samples were randomly distributed across eight LPAs (**Methods, Table S6**). Each color patch represents the arithmetic mean of a single population of cells.

The model accurately predicts the response of CcaSR to all four light signals (**Fig. 4**). “Mono UV” presents the greatest challenge, resulting in an RMSE of 0.15 (**Fig. 4a**). We suspect that prediction errors in this program are due to PCB photodegradation, as we observed no significant toxicity via bacterial growth rate, and the prediction remains accurate until UV reaches maximum intensity (20 µmol m^−2^ s^−1^). “Green mono” (**Fig. 4b**) results in the lowest error (RMSE = 0.038), which is expected because this LED was used to perform the dynamic calibrations (**Fig. 2b,c**). As intended, “Red perturbation” results in an enormous deviation from the reference signal (**Fig. 4c**), and the model accurately predicts this effect (RMSE = 0.081). Finally, “Red compensation” demonstrates that the effect of the perturbation can be eliminated using our model (**Fig. 4d**, RMSE = 0.078).

### Cph8-OmpR photoconversion model

To evaluate the generality of our approach, we repeated the entire workflow for Cph8-OmpR (**Fig. S8–13, Table S5, File S6-7**). Though CcaSR and Cph8-OmpR are both photoreversible TCSs, they have different photosensory domains, ground state activities, and dynamics. To account for the fact that Cph8-OmpR is produced in an active ground state, we used a repressing Hill function (**Methods**). The model again fits exceptionally well to the experimental data (**Fig S8–11**). Unlike CcaSR, which exhibited no detectable dark reversion (**Fig. 2f**), Cph8-OmpR appears to revert in 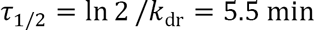 (**Fig. S8f**). As before, *k* is underdetermined, and we chose the best-fit value (**Table S5**). The Cph8-OmpR model performs similarly to its CcaSR counterpart in the spectral validation experiments (**Fig. S12**), and demonstrates greater predictive control in the dynamical validation experiments (**Fig. S13**).

### Development of a CcaSR, Cph8-OmpR dual-system model

We engineered a three-plasmid system (**Fig. S1**) to express CcaSR and Cph8-OmpR in the same cell with sfGFP and mCherry outputs, respectively (**Fig. 5a**). Because the photoconversion parameters are a property of the photoreceptors themselves, we left them unchanged. To recalibrate for mCherry (quantified in Molecules of Equivalent Cy5 (MECY)) and any changes due to the new cellular context, we measured the steady state levels of the sfGFP and mCherry at different combinations of green (*λ*_*c*_ = 526) and red (*λ*_*c*_ = 657) light (**Fig. 5b, S14, File S8**) and refit the Hill function parameters (**Table S6**). The dual-system model accurately captures the experimental observations from the characterization dataset (**Fig. 5b**).

To validate the dual-system model, we again used the biological function generator approach (**Fig. 6**). We designed a series of four dual sfGFP/mCherry expression programs to increasingly challenge the model: “Green mono” using only green light and intended only to control CcaSR (**Fig. 6a**), “Red mono” using only red light and intended to control only Cph8-OmpR (**Fig. 6b**), “Sum”, a simple combination of the first two programs (**Fig. 6c**), and “Compensated sum” where the green light time course is re-optimized to account for the presence of the red signal (**Fig. 6d**) as before (**Methods**). Due to the minimal response of dual-system Cph8-OmpR to green light (**Fig. 5b**), there was no need to adjust the red program to compensate for the presence of green light. The validation experimental results (**Fig. 6**) show that our dual-system model accurately captures both the sfGFP and mCherry expression dynamics. The CcaSR predictions are nearly as accurate as the single-system experiments (**Fig. 4**), and the Cph8-OmpR results match single-system accuracy (**Fig. S12–13**), demonstrating the extensibility of our approach to multiple optogenetic tools.

**Fig 6.**
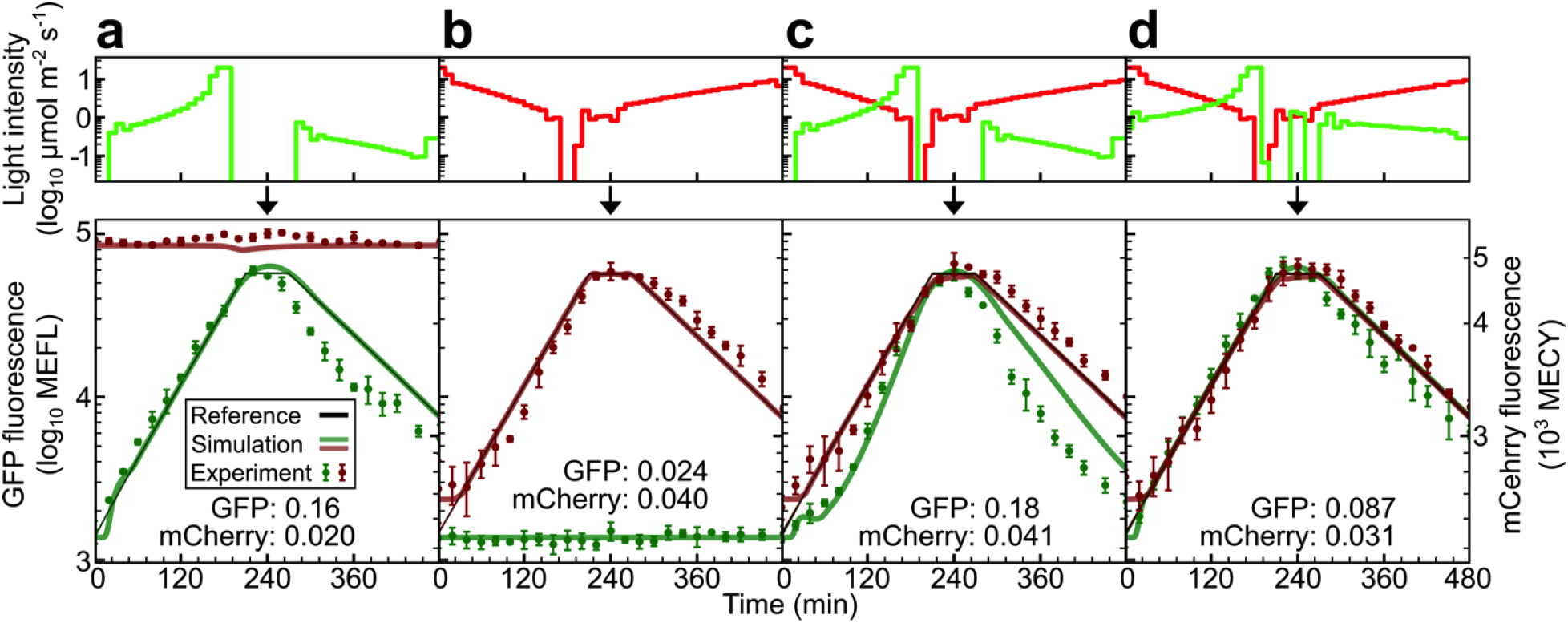
Validation of the multiplexed system model. Predicted responses of the multiplexed system (Fig.5a) to time-varying signals of green (*λ*_*c*_ = 526 nm) and red (*λ*_*c*_ = 657 nm) light are compared to experimental results. Reference signals, light programs, and experimental data are as in **Fig 4**. (a) “Green mono”. The green LED alone provides an optimized input signal for CcaSR. (b) “Red mono”. The red LED alone provides an optimized input for Cph8-OmpR. (c) “Sum”. The “Green mono” and “Red mono” programs are used simultaneously without any compensation, leading to a substantial deviation of the CcaSR output from the reference trajectory. (d) “Compensated sum”. The “Red mono” program is used; however, the green light program is produced while incorporating red light program into the LPG (above). RMSE relative errors are expressed in log_10_ decades (**Methods**). Error bars correspond to the standard deviation in fluorescence measurements over three separate experimental trials (**Table S6**).

### Multiplexed biological function generation

Finally, we designed and experimentally implemented four multiplexed sfGFP/mCherry expression functions representing classes of signals useful for gene circuit characterization (**File S5-6**). “Dual sines” illustrates that two gene expression sinusoids with different offsets, amplitudes, and periods can be composed without interference (**Fig. 7a**). Variations of this combination of signals could be used to perform frequency analysis of multiple nodes in a gene network. “Sine and stairs” demonstrates that our approach can generate two completely different gene expression signals at the same time (**Fig. 7b**). “Dual stairs” demonstrates that the ratio of two proteins can be varied over a remarkably wide range (**Fig. 7c**). Finally, “Time-shifted waveform” (**Fig. 7d**) demonstrates that our approach can be used to characterize genetic circuits where time-delays are critical, such as those involved in cellular decision-making.

**Fig 7.**
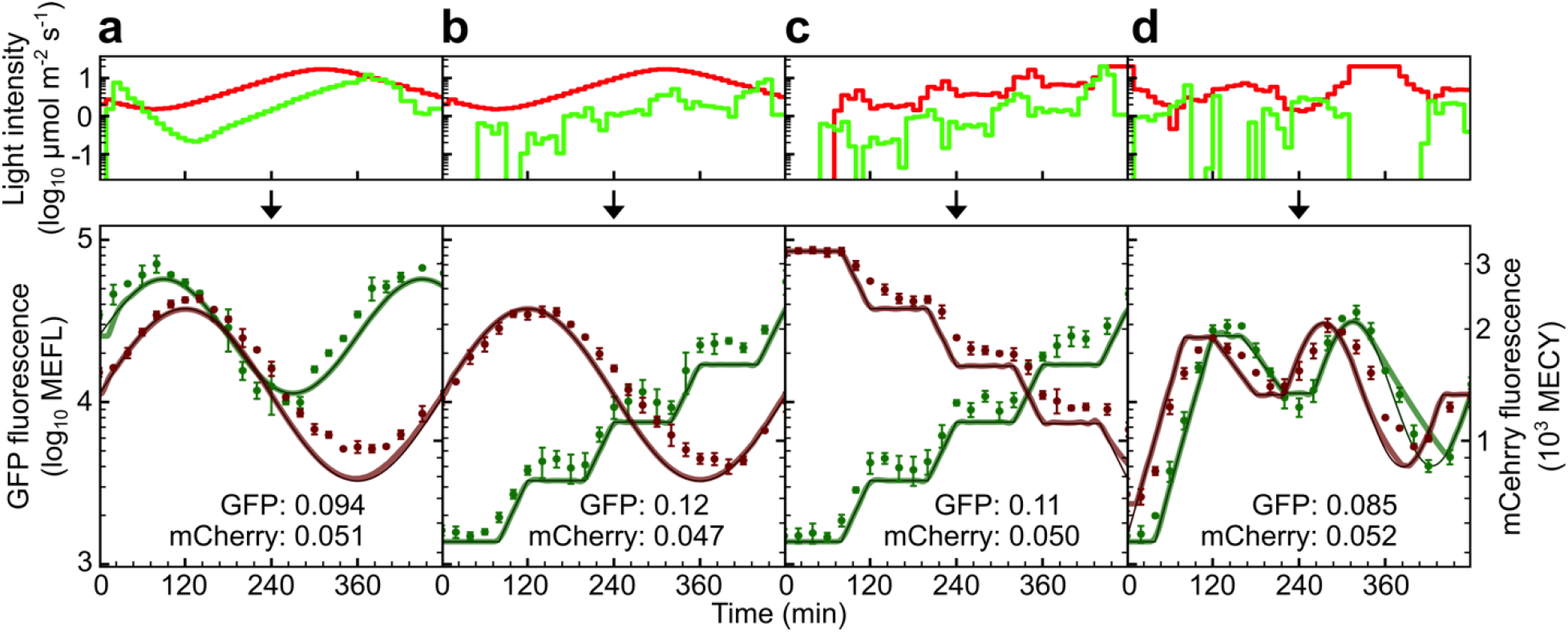
Multiplexed biological function generation. The LPG is used to program CcaSR and Cph8-OmpR outputs to independently follow different reference signals. Red light (*λ*_*c*_ = 657 nm) programs are optimized first using the LPG, and then the “Compensated” approach (**Fig. 6d**) is utilized to generate the green light (*λ*_*c*_ = 526 nm) program (**Methods**). The LPG, reference signals, light programs, and results are as in **Fig 5d**. (a) “Dual-sines”. The sfGFP and mCherry reference trajectories are sinusoids with different periods, amplitudes, and offsets. (b) “Sine and stairs”. The mCherry signal follows the same sinusoid in “Dual-sines,” but the sfGFP reference is a stepped trajectory with several plateaus and increasing linear ramps. (c) “Dual-stairs”. The sfGFP signal follows the same stair-shape in “Sine and stairs,” however the mCherry response is a decreasing stair-shape. (d) “Time-shifted waveform”. The sfGFP and mCherry reference trajectories both follow the same arbitrary waveform consisting of ramps, holds, and a sinusoid, with sfGFP trailing mCherry by 40-minutes. RMSE relative errors are expressed in log_10_ decades (**Methods**). Error bars correspond to the standard deviation in fluorescence measurements over three independent experimental trials (**Table S6**).

## Discussion

In this study, we demonstrate the first use of a mechanistic model of wavelength-dependent photoconversion to characterize and control light responsive signaling pathways *in vivo.* Additionally, we develop a standard set of characterization and validation experiments to parameterize the model and demonstrate that it accurately predicts the spectral and dynamical performance of these optogenetic tools. We demonstrate that the models can be used with virtually any light source or mixture of light sources as long as their emission spectra are known. Finally, we exploit this unique predictive power to demonstrate the first programming of two independent gene expression signals by accounting for inherent cross-talk in the action spectra of the two optogenetic tools that would otherwise impede such efforts.

Our TCS photoconversion model is superior to current alternatives by several key criteria. First, like our previous model^11^, it is quantitatively predictive and requires no parameter recalibrations from day-to-day. However, while our previous model is restricted to a single light source, our current model generalizes to virtually any light source or mixture of light sources. Second, our TCS photoconversion model is compatible with photoreceptors with very different action spectra, opposite ground versus active state signaling logic, and dramatically different dark reversion timescales. Third, our current model modularly decouples the processes of sensing (photoconversion) and output (signal transduction and gene expression). The sensing model component (**Fig. 1a**) should be compatible with a wide range of photoreceptors, including those in other organisms, because the core two-state photoswitching mechanism is used to describe their performance *in vitro*. Then, to describe optogenetic tools based upon those photoreceptors, our TCS output model can be replaced with alternatives appropriate to other pathways, as needed.

A major current problem in optogenetics is that tools developed in different studies are characterized using different culturing conditions, experiments, light sources, reporters, metrics, and so on. This lack of standardization makes it challenging to compare the performance features of different optogenetic tools on even a qualitative basis. The modeling and characterization approach we develop here could be used to make data sheets that describe the behavior of diverse optogenetic tools in standard units. This would enable researchers to choose the most appropriate tool for different applications. Additionally, shortcomings of specific tools could be identified, informing efforts to optimize performance by rational approaches such as protein design^16–18^.

Our approach should enable better control of optogenetic tools with alternative or highly constrained optical hardware used in many research laboratories. For example, many groups perform single cell optogenetic studies using fluorescence microscopes with severely restricted optical configurations. Alternatively, consumer projectors or tablet displays are potentially powerful, low cost hardware options for optogenetics^19,20^. The output spectrum of the light source can be measured and integrated into our workflow. After a simple recalibration (e.g. **Fig. 5**) to account for any changes due to the new growth environment, one should be able to predict and control the optogenetic tool using the new light source.

Oftentimes, it is desirable to simultaneously control an optogenetic tool while imaging a cell of interest using white light sources and excitation light for fluorescent reporters. Such alternative sources of illumination can have deleterious effects on the ability to control the optogenetic tool. However, if the nature of the alternative light signal is known, our approach can compensate for such perturbations (e.g. **Fig. 6, 7**). *In silico* feedback control has also been used to drive desired gene expression dynamics in optogenetic experiments^21–23^. The major benefit of this approach is that perturbations of unknown origin can be compensated by monitoring deviations in the output of an optogenetic tool relative to a reference. Our model is compatible with *in silico* feedback control.

While basic multichromatic control of optogenetic tools has been previously demonstrated^8,24^, the multiplexed biological function generation approach demonstrated here dramatically extends the capabilities of these systems, enabling implementation of several classes of experiments. First, the two-dimensional response of a genetic circuit or signaling pathway could be rapidly evaluated with high reproducibility and precision. For example, one could map the response of 2-input transcriptional logic gates^25^, which integrate the expression levels of two different transcription factors by systematically and independently varying their expression levels while measuring the gate output with a reporter gene. The dynamics of such gates are otherwise difficult to evaluate and seldom characterized^26^. Second, the input/output dynamics of a transcriptional circuit could be characterized as a function of the state of the circuit itself. For example, one could evaluate how well a synthetic transcriptional oscillator can be entrained^27,28^ as a function of the strength of a feedback node. In this case, one optogenetic tool could be used for the entrainment, while the second was used to alter expression level of a circuit transcription factor regulating feedback strength. Third, transcription and proteolysis^29^ could be independently controlled with two different optogenetic tools to alternatively program rapid increases or decreases in expression level. Such an approach could accelerate the gene expression signals that we have generated in this and our previous study^11^, enabling characterization of gene circuit dynamics on faster timescales. Finally, multiplexed biological function generation could be used to evaluate how the timing of expression of two genes impacts cellular decision making^30–32^. For example, in *B. subtilis,* the gene circuits that regulate sporulation and competence compete via a ‘molecular race’ in the levels of the corresponding master regulators^30^. By placing them under independent optogenetic control, the means by which their dynamics impact these cellular decisions could be evaluated more easily and rigorously.

## Materials and Methods

### Bacterial strains

All systems utilize the *E. coli* BW29655 host strain^33^. The CcaSR system strain carries the pSR43.6 and pSR58.6 plasmids, which confer spectinomycin and chloramphenicol resistance, respectively^9^. The Cph8-OmpR system strain carries the pSR33.4 (spectinomycin) and pSR59.4 (ampicillin) plasmids^9^. The dual-system strain carries pSR58.6, pSR78 (spectinomycin), and pSR83 (ampicillin).

### Bacterial growth and light exposure

Cell culturing and harvesting protocols were developed to ensure a high degree of precision and reproducibility in experiments both from well-to-well and from day-to-day (**Note S1**). Cells were grown at 37°C and shaken at 250 rpm throughout the experiment (Sheldon Manufacturing Inc. SI9R) with temperature calibrated and logged by placing a thermometer probe in a sealed 125 mL water-filled flask (Traceable Excursion-Trac 6433). Cultures were grown in M9 media supplemented with 0.2% casamino acids, 0.4% glucose, and appropriate antibiotics. Precultures were prepared in advance by freezing 100 µL aliquots of early exponential phase cultures (OD_600_ = 0.1−0.2) grown in the same media conditions at −80°C (**Note S2**). Cultures were inoculated at low densities (typically OD600 = 1 × 10^−5^) to ensure that final densities did not reach stationary phase (OD600 < 0.2). For each experiment, 192 cultures were grown in 500 µL volumes within 24-well plates (ArcticWhite AWLS−303008), sealed with adhesive foil (VWR 60941–126).

Experiments were performed using eight 24-well Light Plate Apparatus (LPA) instruments^34^, enabling precise control of two LEDs to define the optical environment of 192 cultures at a time. LPA program files were generated using Iris^34^ and Python scripts.

### LED measurement

All LEDs were measured and calibrated (**Note S3**) using a spectrometer (StellarNet UVN-SR-25 LT16) with NIST-traceable factory calibrations performed on both its wavelength and intensity axes immediately prior to use for this study. A six-inch integrating sphere (StellarNet IS6) was used, enabling measurement of the total power output of each LED (in µmol s^−1^). The spectrophotometer was blanked by a measurement of a dark sample before each LED measurement. Measurements were saved as .IRR files, which contain the complete LED spectral power density *P*_light_(*λ*) (µmol s^−1^ nm^−1^) in 0.5 nm increments as well as all setup parameters for the measurement (i.e. integration time and number of scans to average). These files were processed by Python scripts to calculate the LED characteristics, including the peak, centroid, FWHM, and total power. For spectral validation experiments, cinematic lighting filters (Roscolux) were cut, formed into LED-shaped caps, and fitted atop white LEDs (**Table S1**).

### Calculation of *n*_light_

Because the LEDs we utilize have fixed spectral characteristics, the spectral flux density (µmol *m*^−2^ s^−1^ nm^−1^) incident on the photoreceptors can be parameterized by the LED intensity (µmol *m*^−2^ s^−1^). The cultures are shaken throughout the experiment, and we assume that the cells are well mixed within the culture volume. Thus, the mean light intensity within the culture volume, *n*_light_ (*λ*), can be calculated by integrating the intensity throughout the volume of the well. Under the assumption of negligible light absorption by the culture sample (the M9 media is transparent, and the cultures are harvested at low density), this integral simplifies to become the total power of the LED (µmol s^−1^) divided by the cross-sectional area of the well. Given a well radius of 7.5 mm, we calculate 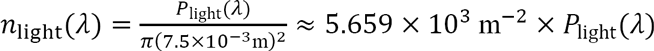.

### LED calibration

Each of the approximately 700 individual LEDs used in the study were measured (**Note S3**), enabling compensation for variation in LED and LPA manufacturing (**Table S1–3**). Each LED was calibrated while powered from the same LPA socket used in experiments. First, a sample of LEDs were measured to identify the electrical current required to achieve an appropriate level of total flux, ∫ *n*_light_(*λ*)dλ. The amount of current required varied depending on the wavelength and manufacturer. The current was adjusted using the LPA ‘dot-correction (DC)’ to achieve a total flux approximately 20% above 20 µmol m^−2^ s^−1^ when the LED was fully illuminated. The appropriate DC level was determined for each LED model. Using these DC levels, the complete set of LEDs were measured. LEDs that produced a total flux below 20 µmol m^−2^ s^−1^ were re-measured at a higher DC level. This set of LED measurements was used to convert the desired intensity time course of each LED into a series of 12-bit grayscale values (i.e. 0–4095) used by the LPA. The LPA reads the grayscale values to produce the appropriate pulse-width-modulated (PWM) signal to achieve the desired intensities.

### Bacterial sample harvesting

Cultures were harvested for measurement (**Note S1**) after precisely 8 h growth by placing the 24-well plates into ice-water baths. Each culture was then subjected to both an absorbance measurement to ensure consistent well-to-well and day-to-day growth, and flow cytometry for quantification of sfGFP or mCherry expression. Absorbance measurements were performed in black-walled, clear-bottomed 96-well plates (VWR 82050-748) in a plate reader (Tecan Infinite M200 Pro). Before fluorescence measurements were performed, culture samples were processed via a fluorescence maturation protocol to ensure measurements were representative of the total amount of produced fluorescent reporter^11^. Rifampicin (Tokyo Chemical Industry R0079), was dissolved in Phosphate-buffered saline (PBS, VWR 72060-035) at 500 µg/mL and used to inhibit sfGFP production during maturation.

### Flow cytometry

Population distributions of fluorescence were measured for each culture on a flow cytometer as previously described^11^. A calibration bead sample (Spherotech RCP-30-5A) in PBS was measured immediately prior to the culture samples from each experimental trial. At least 5,000 events were collected for the calibration bead sample, and at least 20,000 events were collected for each culture sample.

### Flow cytometry data analysis

Single-cell distributions of sfGFP fluorescence were gated, analyzed, and calibrated into MEFL and MECY units using FlowCal^35^. Measurements were gated on the FSC and SSC channels using a gate fraction of 0.3 for calibration beads and 0.8 for cellular samples^35^. Reported culture fluorescence values are the arithmetic means of the cellular populations.

### Sensing model

The light sensing model can be described by the following system of ODEs:

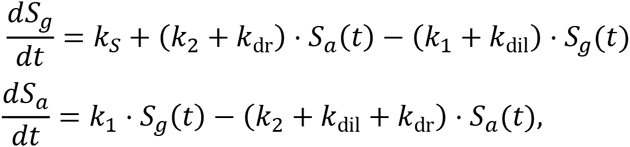

where the variables and rates have been described in the text and figures. Note that *k*_1_ and *k*_2_ are implicitly dependent upon time, as they are functions of the time-varying light environment of the sensors.

If we substitute for the fraction of active sensors, 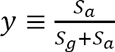, the system reduces to:

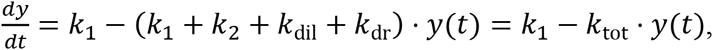

where *k*_tot_ ≡ *k*_1_ + *k*_2_ + *k*_dil_ + *k*_dr_.

This ODE can be solved analytically for a step-change in light from one environment to another. If the step-change occurs at time *t* = 0, then *k*_1_,*k*_*2*_, and *k*_*tot*_ are all fixed for *t* > 0. Given an initial sensor fraction *y*(0) = *y*_0_, we find.

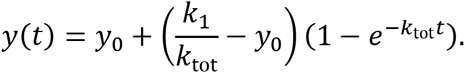

This solution represents an exponential transition from an initial sensor fraction of *y*_0_ to a final fraction given by 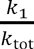 with a time constant set by *k*_tot_. As a result, we anticipate that the transition dynamics of *y*(*t*) will be slowest under zero illumination when *k*_tot_ = *k*_dil_ + *k*_dr_. We also expect that the transition rates will be unbounded as intensity increases.

Finally, for multiple light sources, we simply linearly combine the photoconversion rates from each source: 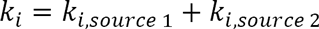.

### TCS signaling model

We utilize a highly simplified model of TCS signaling and gene regulation. This model relates the production rate of the output gene *k*_*G*_(*t*) to the active ratio of light sensors 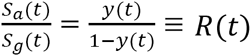. We model TCS signaling as a pure time delay *τ* and a sigmoidal Hill function. For CcaSR, the Hill function is activated by increasing sensor ratios, while for Cph8-OmpR the inverted TCS signaling activity results in a repressing Hill function. Thus, we write 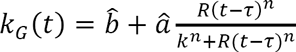 for CcaSR and 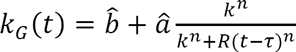 for Cph8-OmpR.

### Output gene expression model

We model output gene expression by first-order production and dilution dynamics:

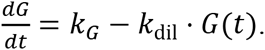

### Generation of model simulations

Simulations were produced by numerically integrating the system of ODEs using Python’s scipy.integrate.ode method using the ‘zvode’ integrator with a maximum of 3000 steps.

### Model parameterization

The CcaSR and Cph8-OmpR models were parameterized using global fits of the model parameters to the complete training data sets (**Fig. 2b-e, Fig. S8b-e**). The ‘lmfit’ Python package, which is based on the Levenberg-Marquardt minimization algorithm, was used to perform the fits and analyze the resulting parameter sets^36^. The fits were performed by minimizing the sum of the square of the relative error between each measured data point and the same point in a corresponding model simulation. Thus the form of the error metric utilized was error 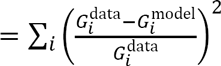 across the complete set of data points 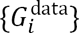.

### Estimation of PCSs

PCS estimates 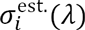 were constructed by linearly regressing a cubic spline to the experimentally determined photoconversion rates in order to produce a continuous PCS (**Fig. S6**). The 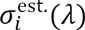 were produced by minimizing the error between unit experimental photoconversion rates 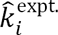 (**Fig. 2f, Fig. S8f**) and spline-derived estimates 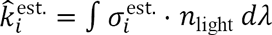. The splines were constructed by establishing a series of integral constraints for the photoconversion rates, continuity constraints for the spline knots, and boundary constraints. As this problem contains more constraints than parameters, optimization is required. We used weighted least-squares with Lagrange multipliers to optimize each spline. To avoid over-parameterization of the 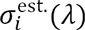, we used “Leave-one-out cross-validation (LOOCV)” to evaluate the performance of optimal splines with between 5 and 20 knots in order to determine the ideal number required for each 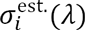 (**Fig. S7**).

### Calculation of prediction error (RMSEs)

For model validation we use a relative error metric 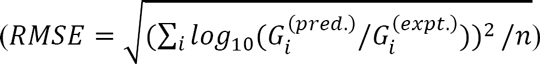 that reports the root-mean-square (RMS) of the log_10_ error between the predicted and measured responses.

### Light program generator (LPG) algorithm

The light program generator was used as previously described^11^. The only modification was to use simulations generated by the model described herein rather than the previous model. Compensated light programs were generated by incorporating the presence of the external light signal into the model simulations.

## Acknowledgements

We thank Sebastian Schmidl for providing the dual-system strain, Prabha Ramakrishnan and Karl Gerhardt for helpful discussions on the design of the characterization experiments, Sebastian Castillo-Hair for helpful ideas on constructing the photoconversion cross-section estimates, and Keshav Rao for assistance with data collection. This work was supported by the Office of Naval Research (MURI N000141310074) and an NSF CAREER award (1553317).

## Author contributions

EJO and JJT conceived of the project. EJO designed and performed experiments and analyzed data. CNT assisted with the photoconversion cross-section estimates and design/performance of spectral validation trials. EJO and JJT wrote the manuscript.

## SI Materials and Methods

### Detailed bacterial growth and light exposure protocol

#### 1. Experiment initialization. This an approximately 2-hour process

(a) Prepare program files on SD cards and load the SD cards into LPAs.

(b) Prepare 24-well plates for use (ArcticWhite AWLS-303008): Soak plates for at least 15 minutes in 70% EtOH. Triple-rinse the plates using DI water. Place rinsed plates onto a sheet of foil which will later wrap around the plate. Dry plates near a lit burner until the interior of the wells are dry (approximately 45 min). Use a paper wipe to dry the bottom of the plate and the foil. Wrap the plate in the foil.

(c) Prepare media and cell culture: Prepare 100 mL M9 medium (75.8 mL autoclaved, distilled H2O, 20 mL 5x M9 salts, 2mL 10% casamino acids, 2mL 20% glucose, 200 µL 1m MgSO4, 10 µL 1m CaCl2). (Note: prepare 1-2L of this media and pipet it into 50mL aliquots to be used at a later date. For 8x LPA experiments, combine two aliquots.). Add appropriate antibiotics to medium. Shake/stir the container to homogenize. Remove −80 °C culture aliquot from the freezer and allow to thaw. Lightly vortex thawed culture aliquot and briefly spin down in a microcentrifuge. Add appropriate culture volume to the 100mL media. (Note: inoculation densities for strains used in this work are CcaSR: 1E-5, Cph8-OmpR: 5E-6, Dual-system: 1E-5).

(d) Distribute inoculated media into 24-well plates: Prepare a row of 1mL tips in a box so that only every-other tip is present (i.e. 6 tips total). Shake to homogenize inoculated media. Arrange the 8x 24-well plates with the wells open and uncovered. Pour more than half of the media into a 50mL disposable multichannel tray. Use a 1mL 12-channel pipettor with the previously-prepared row of 6 tips to transfer 500 µL of media into each well. Pour remaining media into the tray and continue. Cover each plate with an adhesive-backed foil seal (VWR 60941-126).

(e) Load the LPAs and start the programs: Carry the 8x plates, 8x LPA lids, and 32x LPA wing nuts to the incubator containing the LPAs. (Note: a small carboard box or plastic container is helpful). Load the plates onto each LPA, ensuring that the plate is oriented correctly and is completely engaged with the LPA plate adapter. Place the lid onto each LPA, ensuring that the lid is oriented correctly. Engage 4x LPA wing nuts onto each device. Tighten the nuts evenly until the pressure of the rubber gaskets being compressed is felt (approximately 1-2 full turns after the nut engages with the lid). Start an 8-hour timer while synchronously releasing the reset button on one of the LPAs. Every 10 seconds, release the reset button on another LPA. Make sure to recall the order in which the LPAs were reset. This staggering of the start times enables the plates to be removed immediately when each of their programs ends. Allow the cells to grow in the LPAs at 37 °C for 8 hours.

#### 2. Experiment completion and data collection. This is an approximately 4-hour process

(a) 30 minutes prior to the end of the LPA programs, prepare a PBS+rifampicin solution: Fill a 250 mL beaker with 125 mL of a PBS solution in the pH7-7.2 range (VWR 72060-035). Add a stir-bar to the beaker and place on a stir-plate. Weigh 62.5 mg of rifampicin (rif, Tokyo Chemical Industry, R0079). Adjust the beaker to be centered on the stir plate. Adjust the rate of the stir plate to produce a vortex which doesn’t quite reach down to the bar. Add the rifampicin to the middle of the vortex. Slide the beaker to be slightly off-center on the stir plate. This often encourages the vortex to lower, leading to the stir-bar clipping the vortex with each rotation. This is the desired state for dissolving. Make any adjustments if necessary. Cover the dissolving solution with an opaque container (e.g. tin can) or foil, as the rif is light-sensitive.

(b) Finishing the LPA programs: Prepare two autoclave trays with ice-water baths. Make sure the water level is near the surface of the ice so that the submerged items will make contact with water and not just ice (Note: a utility cart is useful here). Submerge the 8x multichannel-ready tube racks into one of the icewater baths. Load the 192x wells in these racks with flow cytometry tubes (VWR 60818-419). Carry the other ice-water bath to the incubator with the LPAs. At 2 min time remaining on the LPA programs, open the incubator and unscrew all of the wing nuts (leave the lids in place). As the status LEDs on each LPA change to indicate that the program is complete, remove the enclosed plate and immediately submerge it at least halfway up the side of the plate in the ice-water bath. The temperature logs from the data-logging thermometer on the growth incubator can now be gathered.

(c) Preparing for culture transfers: After a 10 minute wait, remove the rif solution from the stir-plate. The solution should be a vivid, bright orange. If it is dark-colored, either there was light leakage, the stirring was too vigorous, or the saline was not in the pH 7-7.2 range, and the solution should be remade. Filter the rif solution using a 0.22 µm filter. Use a multichannel pipettor to load the 192x cytometry tubes each with 500 µL of the rif solution. Label 2x black-walled clear-bottomed 96-well plates (VWR 82050-748). Prepare 192x 200 µL tips by arranging them in four tip boxes such that every-other tip is present (i.e. rows of 6 tips).

(d) Transferring cultures for measurement: Remove the first 24-well plate from its water bath and place it on a paper towel. Use another towel to dry its top surface and sides. Carefully remove and discard the adhesive foil without spilling the contents of the plate. Use a 12-channel 200 µL pipettor to transfer 100 µL volumes of the cultures into the 96-well plate. Make sure to pipette up-and-down in the 24-well plate in several locations to ensure that the culture is homogenized before this transfer. Do not discard the tips after this transfer. After the transfer to the 96-well plate, using the same tips, transfer the same volume into the corresponding PBS+rif tubes in the tube racks. Discard the tips after this transfer. Place plastic caps onto the PBS+rif tubes containing the cultures. This aids in reducing pipetting errors. Repeat this process until all 192x samples have been transferred to the 96-well plates and the cytometry tubes. It is useful to fill the 96-well plates using an interleaved strategy, where the first 24-well plate begins in A1, the second plate in A2, the third plate in E1, and the fourth plate in E2. Once all samples have been transferred, the cytometry tubes should be homogenized. To do this, take the racks out one-at-a-time, tilt them nearly horizontally, and gently shake the rack. Rotate the rack 180° and repeat. This will cause the PBS+rif to roll up the sides of the tubes, mixing in any culture which was stuck on the sides of the tube.

(e) Preparing for the next experiment (optional): If another experiment is to be started as soon as possible (the 8x LPA growth protocol can be performed twice in a day), the 24-well plates should be cleaned immediately and dried after at least a 15-minute soak in 70% ethanol in water.

(f) Fluorescence maturation and culture OD measurements: Transfer the PBS+rif+culture tube racks into a 37 °C water bath. Allow 1 hour for maturation of fluorescent proteins. While the maturation is in process, use a plate reader to measure the absorbance of the cultures in the 96-well plates. When the fluorescence maturation is complete, remove the tube racks and place them back into an ice-water bath. The samples should remain on ice at least 30 minutes before measurement via flow cytometry. The temperature logs from the data-logging thermometer on the maturation incubator can now be gathered.

(g) Calibrated cytometry measurements: Prepare a bead sample for cytometry calibration. Pipette 500 µL of PBS into a cytometry tube. Add one drop of calibration beads (Spherotech RCP-30-5A) to the PBS. Cap the tube and place it on ice. Transport the PBS+rif+culture samples and the bead sample to the cytometer (modified BD FACScan^1^). Turn the cytometer on and allow it to pressurize and stabilize the laser. If applicable to your cytometer, perform several “Drain/Fill” cycles to clear the lines of any residue leached from the tubing. Also fill and drain the sheath and waste containers if necessary. Analyze the bead sample. The beads should be tightly-clustered in the center of the FSC vs. SSC scatter plot. The gain settings on the fluorescence channels should match the gain used to acquire the cell samples (if measuring a strain for the first time, it is wise to measure the cultures anticipated to have the highest and lowest fluorescence levels to establish an appropriate gain). Collect at least 5,000 events (more is better). Run the cell samples through the cytometer. The cells should be clearly visible on the FSC vs. SSC scatter plot. Collect at least 20,000 events per sample. Make sure that the fluorescence histograms are fully contained within the measurement range at the current gain settings. When complete, perform an instrument shutdown cycle by running 10% bleach and then water through both the droplet-containment line and the sample line.

### Detailed −80 °C preculture aliquot protocol

1. Perform a plasmid transformation to produce your desired strain. From the transformation plate, pick a colony and grow a 3 mL culture in LB media + appropriate antibiotics. Grow to a OD600 of 0.1-0.2. Make a standard 1mL −80 °C glycerol stock from this culture (700 µL culture and 300 µL of 60% glycerol) and freeze the stock overnight.

2. Use a sterile toothpick or pipette tip to stab the glycerol stock and begin growing an 8 mL liquid preculture in a culture tube under experimental growth conditions (i.e. 37 °C for the experiments in this manuscript). It is critical to grow this culture using the same media which is specified for the experiments you will be performing with the aliquots. That is, if your experiments will be in M9+antibiotics, you should grow this culture in M9+antibiotics. Prepare a PCR rack with 48 sterile, opened PCR tubes. Cover the tubes with foil.

3. When the culture begins to exhibit the slightest bit of turbidity, take a 1 mL sample from it and check the density. Check the density again in 30 minutes. Use these measurements to (exponentially) extrapolate the time at which the culture will reach an OD600 of 0.1. If the culture OD600 is already above 0.1, but below 0.2, remove immediately. If the culture is above 0.2, start over with a new preculture.

4. When an OD600 of 0.1-0.2 is reached, remove the tube from the shaker. Do not put the culture on ice. Add a quantity of 60% glycerol to the preculture to bring the final glycerol concentration to 18%. Homogenize the culture by gently vortexing. Use a multichannel pipettor to transfer 100 µL of the preculture into each of the PCR tubes. Close all of the PCR tubes, and transfer the rack to a −80 °C freezer.

5. After 30 minutes, transfer the frozen PCR aliquots into a labeled 50 mL conical tube. Store the conical in the −80 °C freezer.

### Detailed LED measurement protocol

1. Initialize the LPA by programming it with your desired settings. For LED calibration, first maximize all of the parameters for each LED by setting the DC values to 63, GCAL values to 255, and GS sequence to 4095. If a specific final intensity is desired, adjust the DC parameter to produce a measurement which is approximately 10-20% above your desired intensity.

2. Connect all wires to the spectrometer (StellarNet UVN-SR-25 LT16): power to wall-socket, USB to computer, and fiber-optic to the IS6.

3. Open the SpectraWiz software and say “No” to the pop-up dialog named “Confirm” if it shows up. If the software has not been installed, follow the manufacturer instructions for installation.

4. Load the IS6 calibration file. Click “View” → “Radiometer” → “Save or Load Cal File for attached light receptor !” and then click “Yes” to open a file explorer dialog. Select the calibration file (“MyCal-EPP10022524-RAD-IS6.CAL” for our instrument) and click “Open.” Click “OK.” at the dialog that pops up.

5. Switch the instrument to scope mode, which reads out the raw 16-bit values from the light detector. Click “View” → “Scope Mode.”

6. Place the LED into the IS6 LED adapter socket, and screw the adapter onto the IS6. The IS6 LED adapter is the screw-on cap with a wire coming out of the back of it terminated by a barrel-jack power connector. Connect power to the barrel-jack connector using the LPA-side of the power adapter. The LPA-side of the power adapter is the wire terminated in a male barrel-jack on one end and a two-pronged terminal on the other. The two-pronged end fits into the LED socket in the LPA in which the currently-measured LED will be placed. Note: to minimize measurement of back-emission from the LED, a 3d-printable cone which holds the LED in place is available.

7. Adjust the integration time on the spectrometer to maximize the signal from the LED. Click “Setup” → “Detector integration time” and type a value which produces a signal that peaks at approximately 75% of the detector range. Repeat if necessary until the full detector range is utilized. Note: By default the software will be averaging the spectra of 5 independent measurements. To speed up the measurement process, the number of repeats can be reduced by clicking “Setup” → “Number of scans to average” and specifying a smaller number.

8. Switch to a calibrated intensity y-axis. Click “View” → “Radiometer” → “MicroMoles per square meter per second” and click “Yes”. You may want to adjust the limits of the y-axis so that the spectrum is visible. To do this, click “View” → “Y scale” → “set Max Y” and specify a lower value. A typical value is 0.0001.

9. Blank the spectrometer. Remove power from the LED by disconnecting the power adapter at the barrel-jack connection. Allow the spectral measurement to update until the spectrum has cleared the screen. Then click the dark light bulb icon (fifth icon from the left near the top of the screen). Reconnect power to the LED and allow the spectrum to stabilize.

10. Export the spectrum. Click the floppy disk icon (second icon from the left near the top of the screen). Choose a location and save the spectrum. If the “File save EXPORT parameters” dialog pops up, click “auto-set” on each of the fields to maximize the exported range and resolution of the spectrum.

11. To measure the next LED, remove the current LED and return to either step 5 or 6. Use step 6 if the next LED has an intensity known to be similar to the just-measured LED, or use step 5 to adjust the integration time if necessary. The output of the LED measurement protocol is a “.IRR” spectrum file which contains the complete LED power spectrum in ∆λ = 0.5nm increments as well as all setup parameters for the measurement (i.e. integration time, number of averages, etc.). This spectra file can be viewed in a text editor or can be loaded into a spreadsheet viewer. The LED power spectrum can be used to calculate a number of characteristics, including the peak, centroid, FWHM, and total flux.

### Comparison of output gene expression ranges for single- vs dual-systems

The CcaSR output range is nearly conserved (60-fold vs. 56-fold) while the mCherry response from Cph8-OmpR is substantially reduced (210-fold vs. 6.0-fold). Additionally, the light response is less sensitive than was observed for Cph8-OmpR individually, as half-repression requires a 5.2-fold higher intensity (**Fig. S16**). We speculate that the reduction in the output range and decrease in sensitivity of Cph8-OmpR results from a competition between Cph8 and CcaS for limiting PCB, leading to a substantial population of light-insensitive apo-Cph8. Notably, the growth rate (**Fig. S15**) of the dual-system strain (39.2 minutes per doubling) is only marginally slower than the single-system strains (37.4 and 37.9 minutes for CcaSR and Cph8-OmpR, respectively).

### Detailed descriptions of multiplexed function generation reference signals

In the below descriptions, the percentages and fractions correspond to a log-scaled representation of the output range (e.g. if a system has a 16-fold output range, the 50% level on a log scale would be at the same expression at the 25% level on a linear scale).

1. Dual-sines. The mCherry reference signal is described by the function 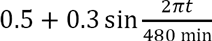 while the sfGFP reference signal follows 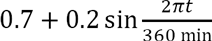
2. Sine and steps. The mCherry reference signal is the same as in “Dual sines”, while the sfGFP signal is a series of 80-minute holds and 40-minute linear ramps in increasing increments of 20% of the output range.
3. The sfGFP signal is the same as in the “Sine and steps” program, while the mCherry signal is the inverse of the same program.
4. (Time-shifted waveform). The mCherry signal is a complex function consisting of
  a. a linear ramp from 0 to 70% over 80 minutes,
  b. a hold at 70% for 40 minutes,
  c. a linear ramp down to 50% over 60 minutes,
  d. a hold at 50% for 40 minutes,
  e. a sine wave described by the function 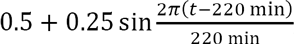
  f. and a hold at 50% for 40 minutes.

The sfGFP signal is the same program but delayed by 60 minutes.

**Fig S1.**
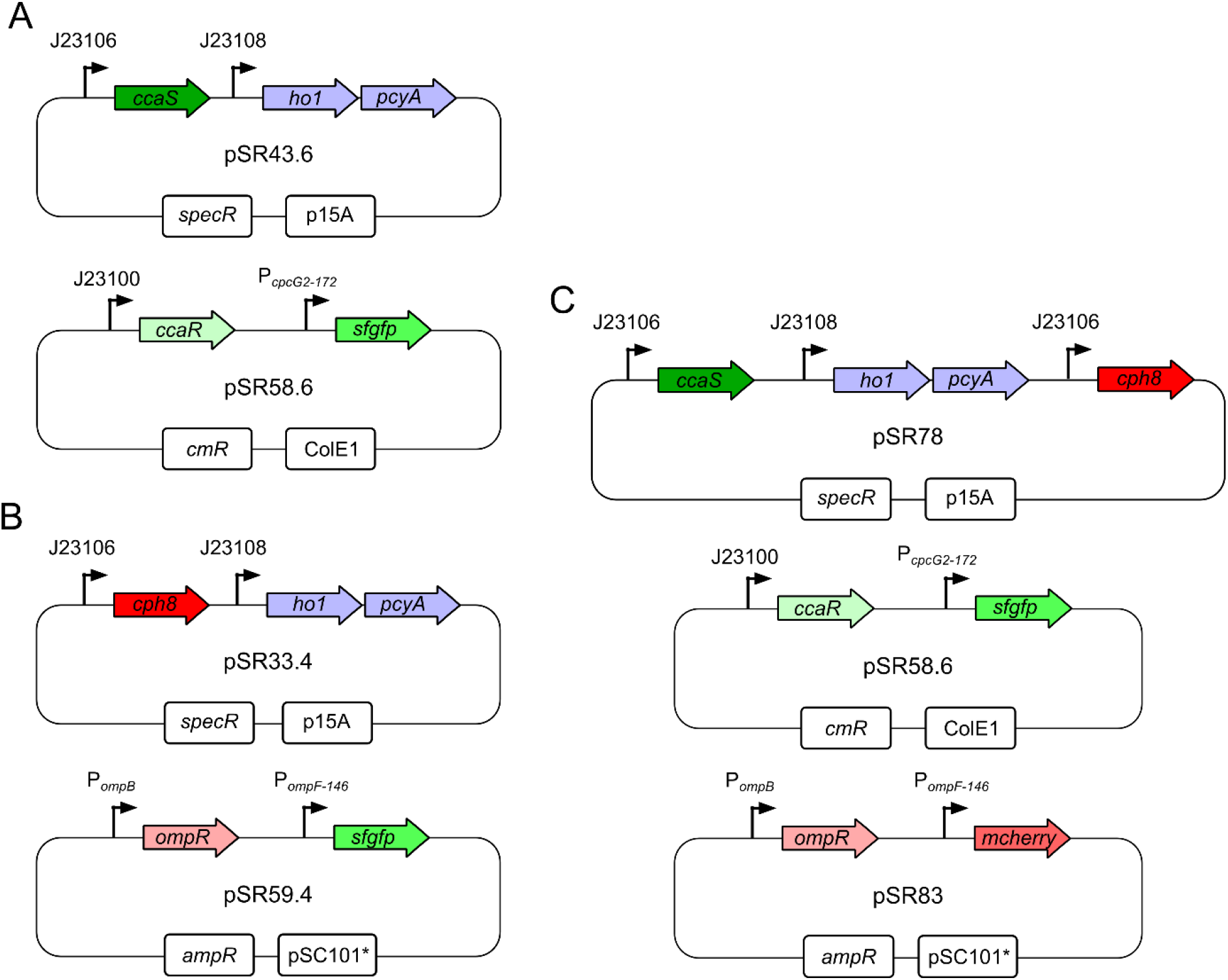
CcaSR, Cph8-OmpR, and dual-system plasmid maps. (A) Maps of plasmids pSR43.6 (top) and pSR58.6 (bottom) expressing the CcaSR system including the PCB biosynthetic enzymes ho1 and pcyA^2^. (B) Maps of plasmids pSR33.4 (top) and pSR59.4 (bottom) expressing the Cph8-OmpR system. (C) Maps of plasmids pSR78 (top), pSR58.6 (middle), and pSR83 (bottom) used in the dual-system experiments.

**Fig S2.**
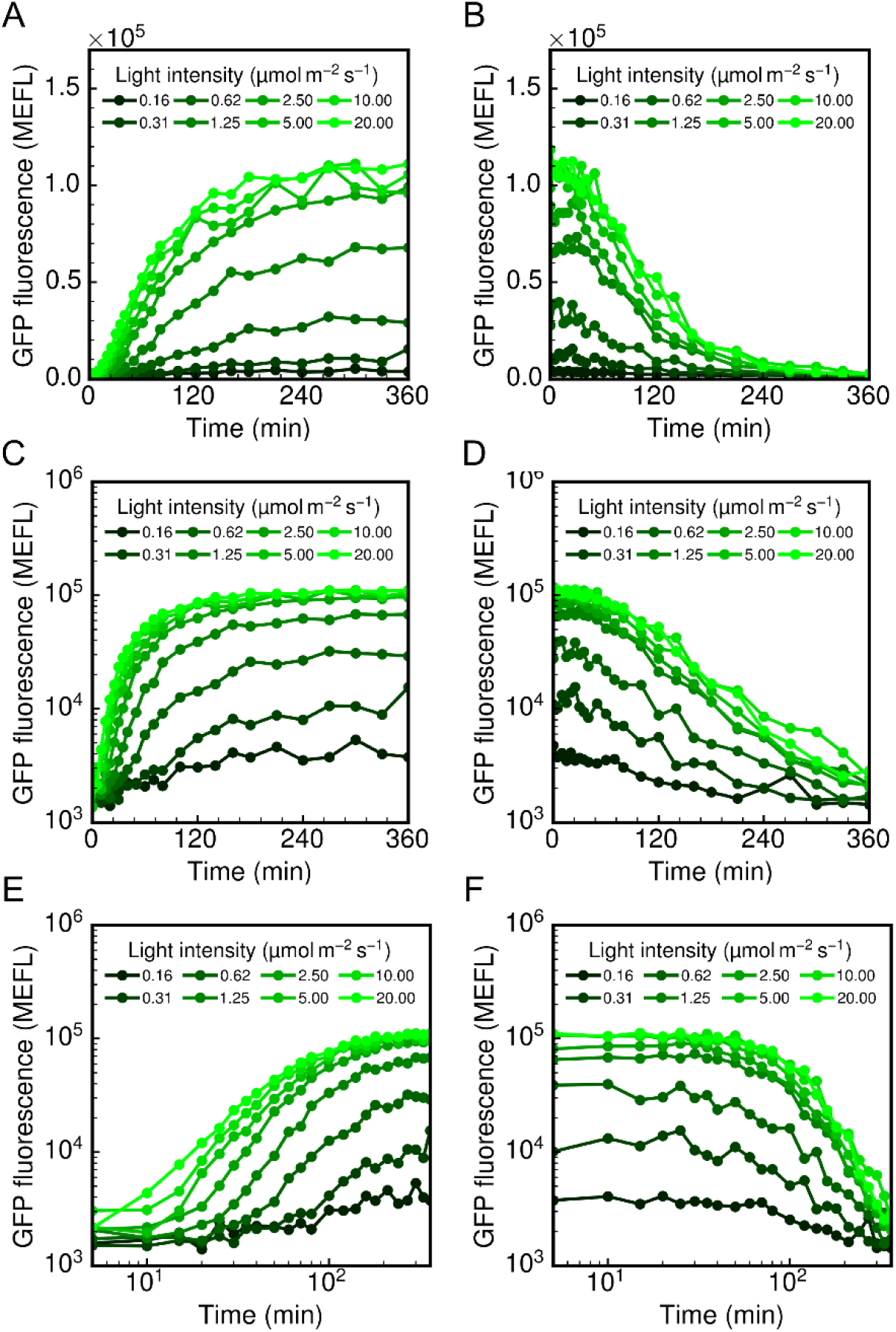
Alternative representations of CcaSR dynamic training experiments. (A,C,E) Activating and (B,D,F) deactivating step response dynamics are shown with (A,B) linear axes, (C,D) semilog axes, and (E,F) log-log axes. Each data point represents the arithmetic mean of a single population of cells. Lines are linear interpolations between points, and are simply a guide to the eye.

**Fig S3.**
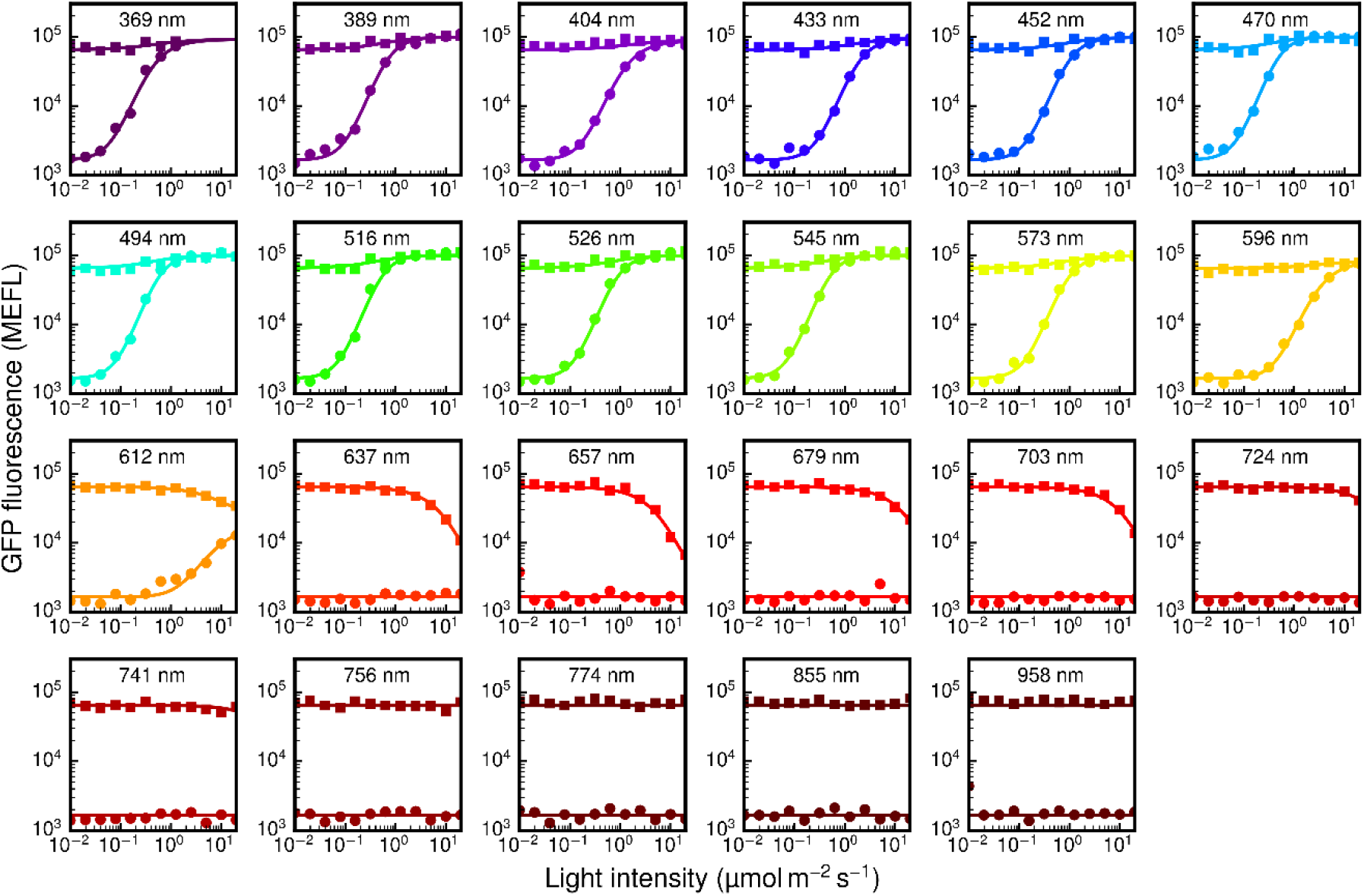
Alternative representations of CcaSR spectral training experiments. Results of the spectral characterization experiments are shown LED-by-LED on log-log axes. Each plot shows the forward activation spectrum (circles) and reverse activation spectrum (squares) for each LED (centroid wavelength indicated). Each data point represents the arithmetic mean of a single population of cells. Lines are simulated results of the best-fit model.

**Fig S4.**
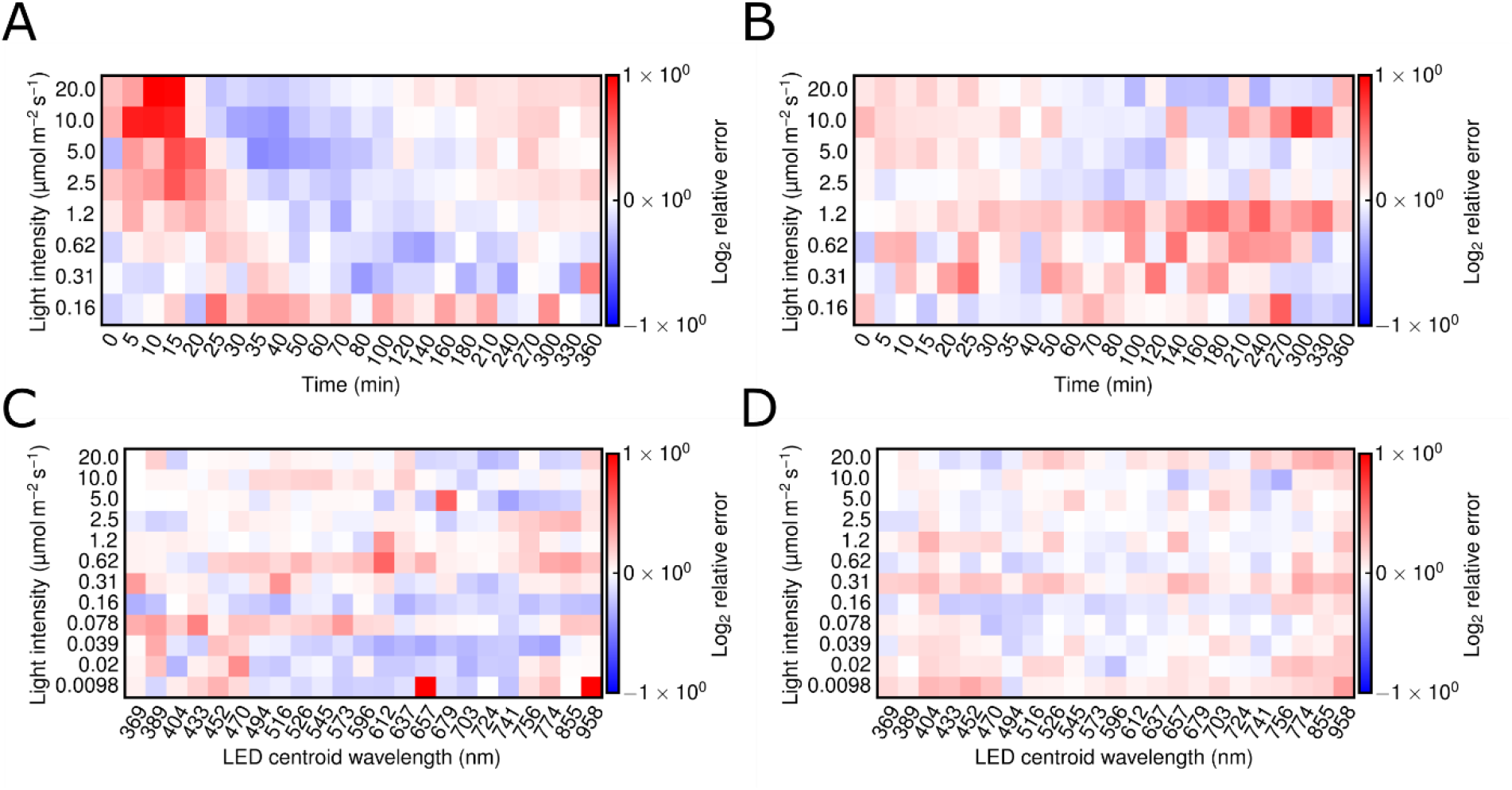
Residuals of CcaSR model to the training data. Residuals between the data and the model are shown for the (A) activating and (B) deactivating step-responses as well as the (C) forward and (D) reverse spectral measurements. The residuals are expressed relative to the measured fluorescence data on a log (base 2) scale (i.e. log_2_ *F*_data_/*F*_model_.

**Fig S5.**
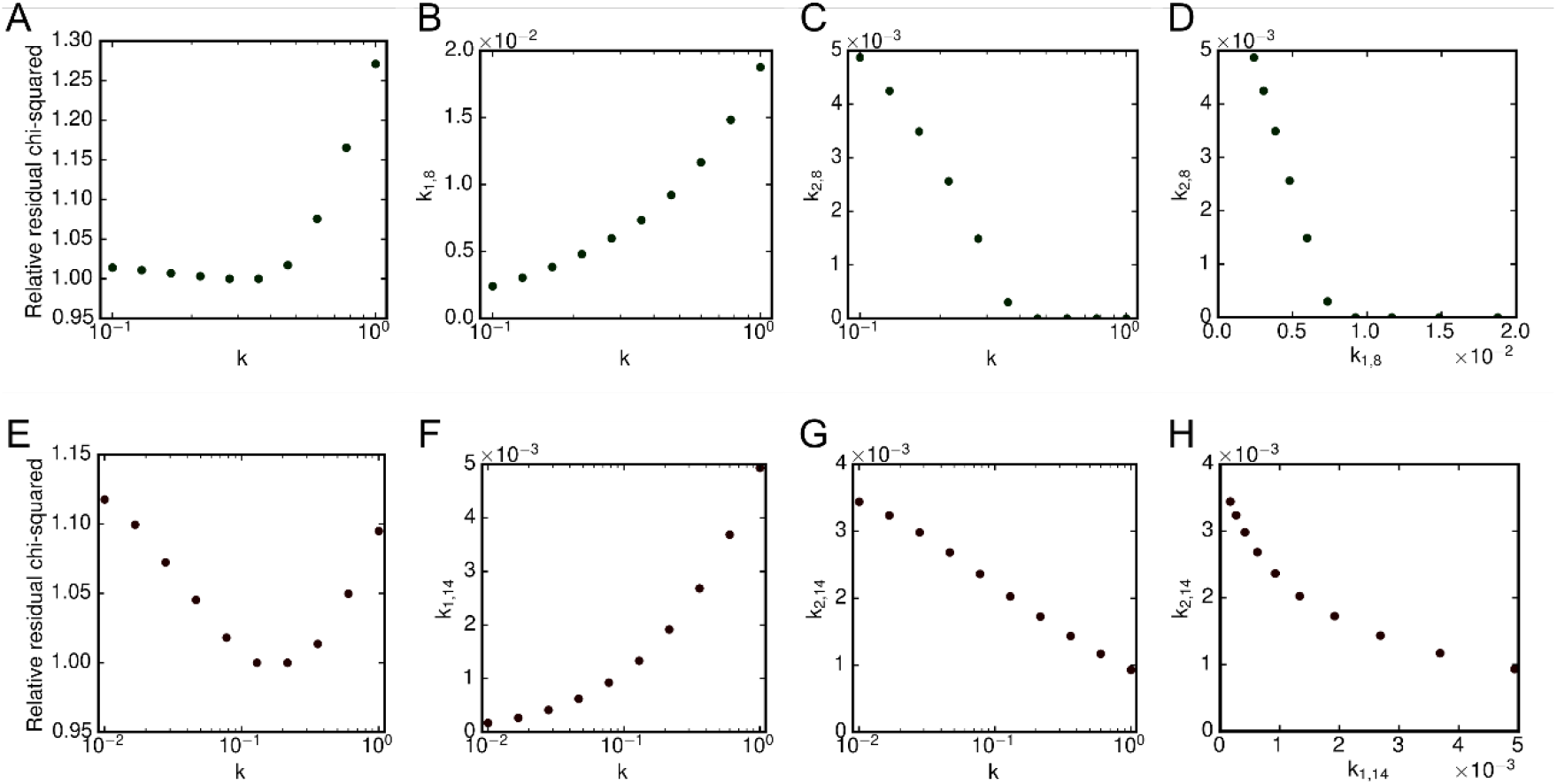
Multicollinearity in the model fit results. The ability of the model parameters to compensate for one another was examined by repeatedly performing the model regression to the data while holding the *k* parameter fixed as an independent variable to a range of values. The CcaSR (A-D) and Cph8-OmpR (E-H) results are shown. The The quality of the fit relative to the best-fit models are shown (A,E), as well as the relationship between the forward (B,F) and reverse (C,G) photoconversion rates. For the CcaSR system, the photoconversion rates for the green LED (*λ*_*C*_=526 nm, *k*_*i,8*_) are shown, and for the Cph8-OmpR system, the red (*λ*_*C*_=657 nm, *k*_*i*,14_) is used. Finally, the forward and reverse rates are plotted against each other (D,H).

**Fig S6.**
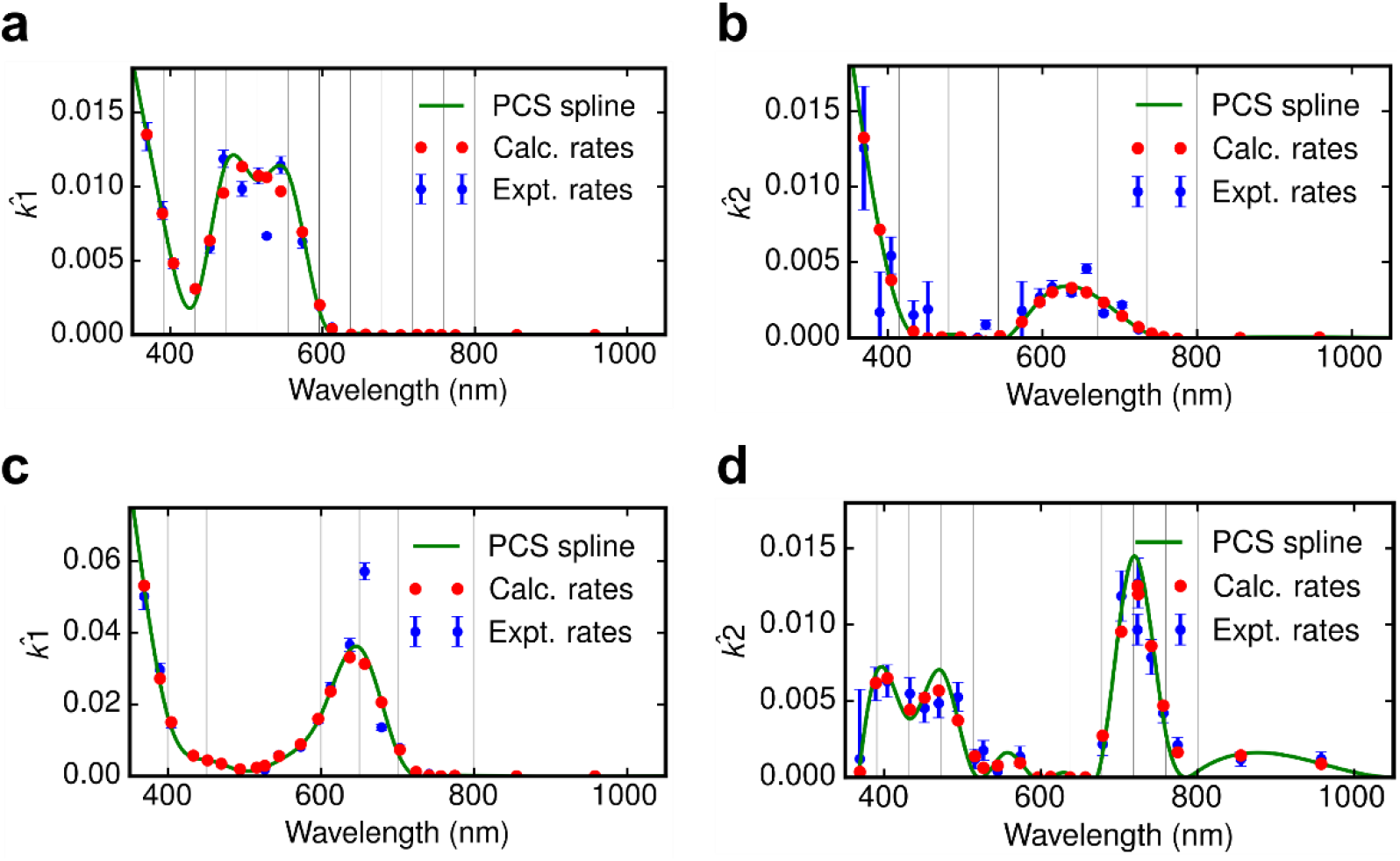
Detailed results of best-fit splines. Spline results and photoconversion rates are shown for CcaSR (a,b) and Cph8-OmpR (c,d). Experimental unit photoconversion rates (blue, 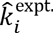) are shown with errorbars indicating the uncertainty via the standard error of the model fits. The PCS spline estimate (green, 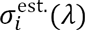) is constructed by minimizing the sum of the squared error (weighted by the experimental uncertainty) between the experimental and spline-derived (calculated) rates. The spline-derived, estimated rates are calculated via 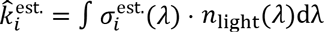. The *λ*_*c*_=526 nm LED was not included during spline optimization, as the experimental rates for this LED appear to be outliers. We believe that because this LED has a wider output angle than the other LEDs, that light from this LED may have been cropped by the LPA well geometry, leading to a lower-than-expected amount of light reaching the cells (and hence lower measured photoconversion rates).

**Fig S7.**
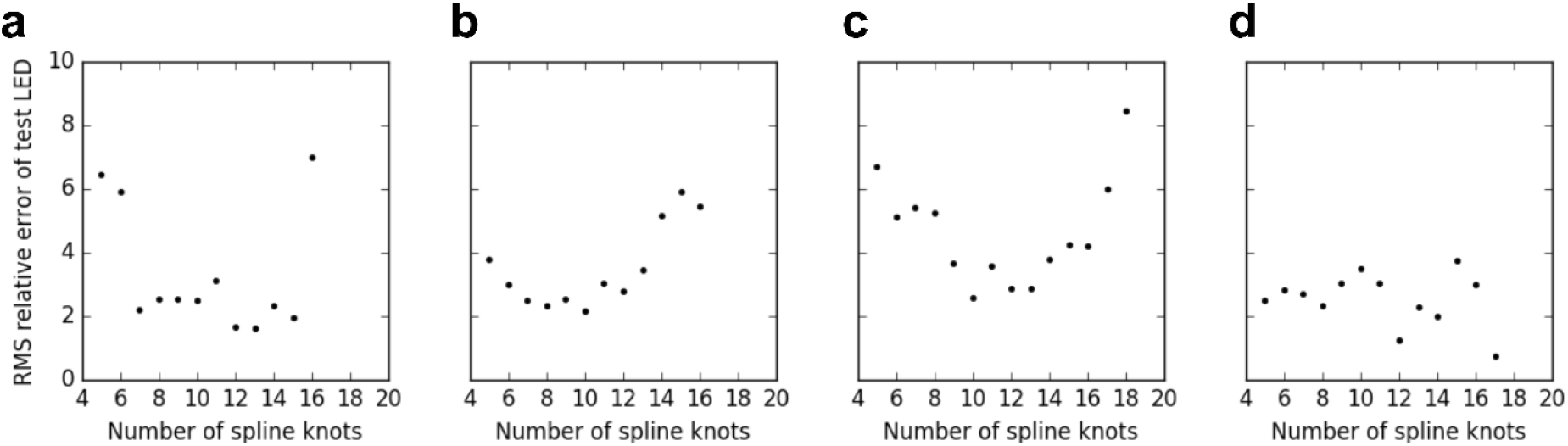
Spline knot number optimization via LOOCV. To determine the appropriate spline complexity (i.e. number of knots) for each PCS estimate spline, we used a “Leave-one-out cross-validation (LOOCV)” approach. In this approach, the experimental dataset is split into a series of training and test LED datasets, each of which is constructed by pulling a single LED out of the training set to be used for testing. Then, for each number of spline knots to evaluate, we construct a spline to each training set (the spline knots are evenly distributed from 350 to 800 nm), predict the remaining test LED, and calculate the RMS of the relative errors of the predictions. The RMS errors for each level of spline complexity are shown for CcaSR forward (a) and reverse (b) spectra and for Cph8-OmpR forward (c) and reverse (d) spectra.

**Fig S8.**
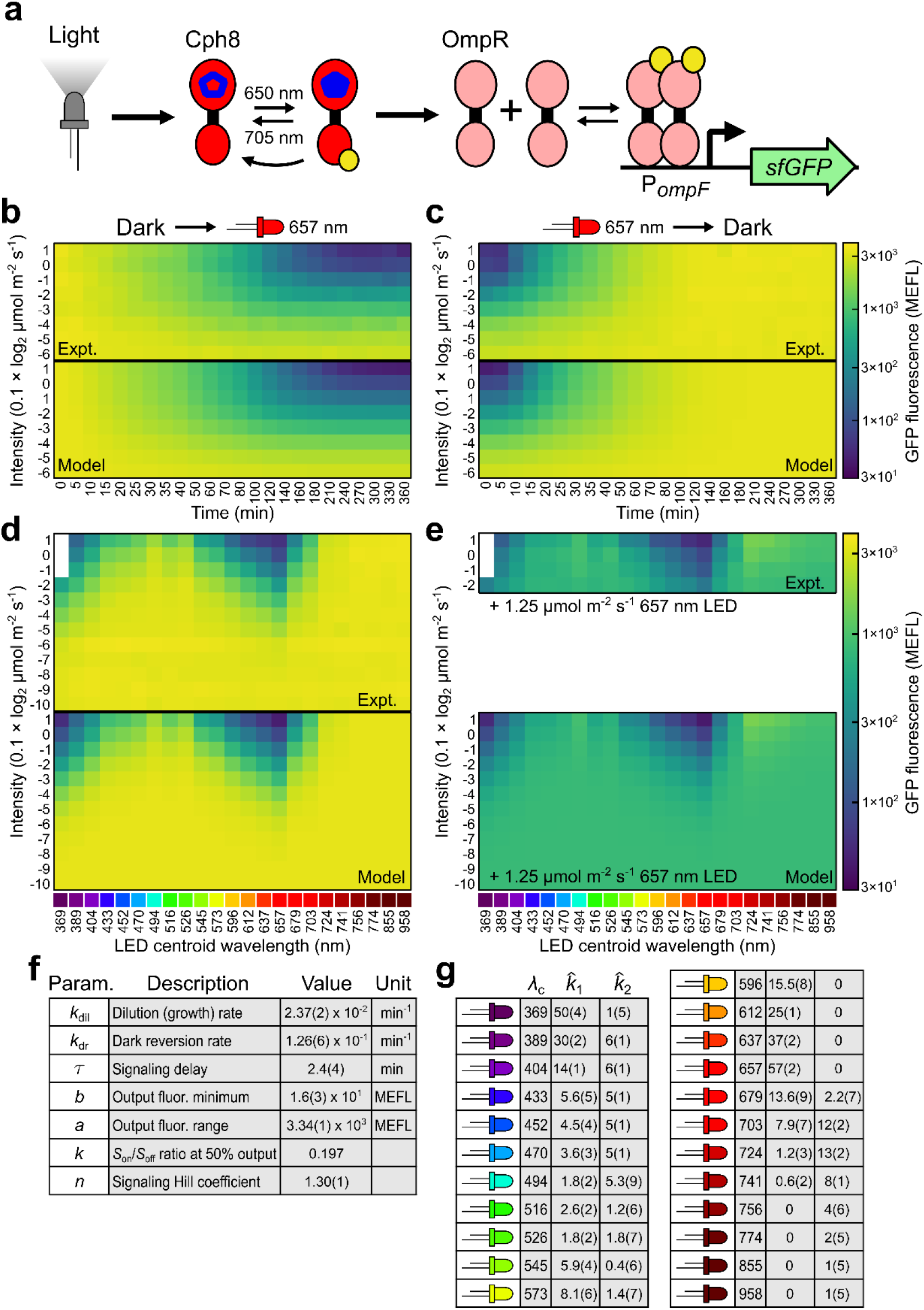
Cph8-OmpR characterization and model parameterization. Figure details are described in Fig. 2. Note that the reverse action spectrum measurements (e) contain only four intensities of the spectral LEDs rather than the full set of 12 used in the CcaSR experiments.

**Fig S9.**
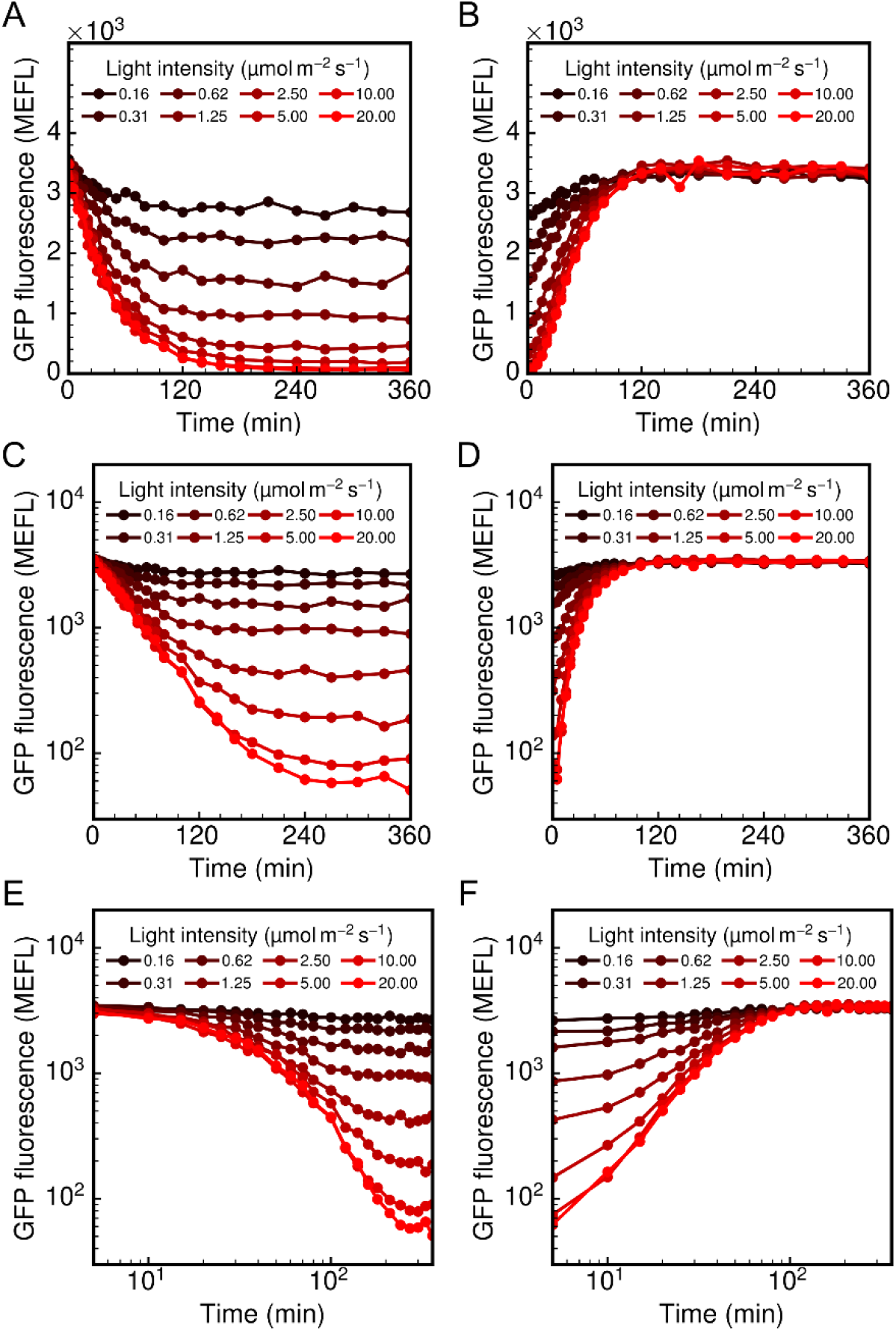
Alternative representations of Cph8-OmpR dynamic training experiments. A,C,E) Activating and (B,D,F) deactivating step response dynamics are shown with (A,B) linear axes, (C,D) semilog axes, and (E,F) log-log axes. Each data point represents the arithmetic mean of a single population of cells. Lines are linear interpolations between points, and are simply a guide to the eye.

**Fig S10.**
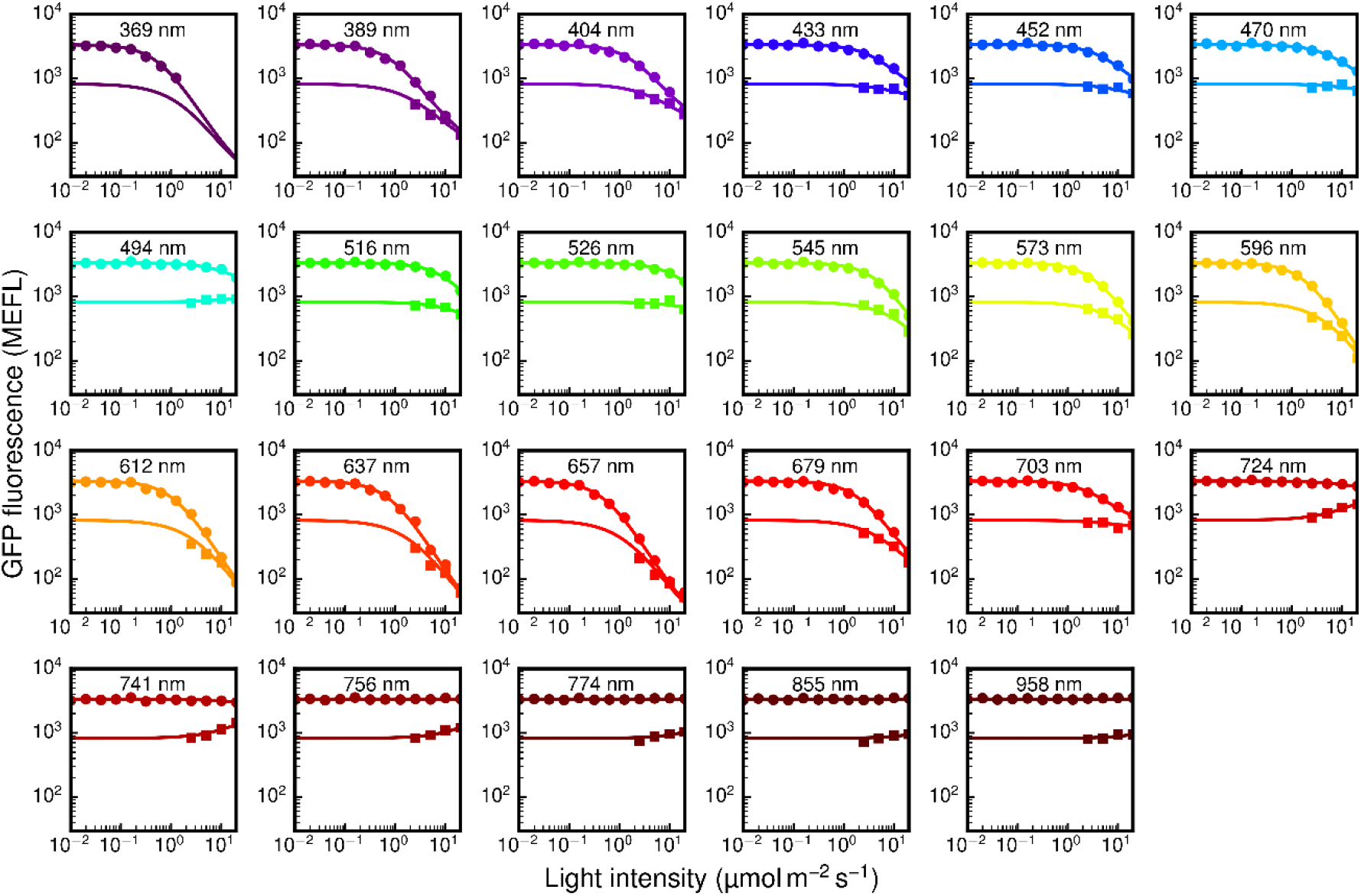
Alternative representations of Cph8-OmpR spectral training experiments. Results of the spectral characterization experiments are shown LED-by-LED on log-log axes. Each plot shows the forward activation spectrum (circles) and reverse activation spectrum (squares) for each LED (centroid wavelength indicated). Each data point represents the arithmetic mean of a single population of cells. Lines are simulated results of the best-fit model.

**Fig S11.**
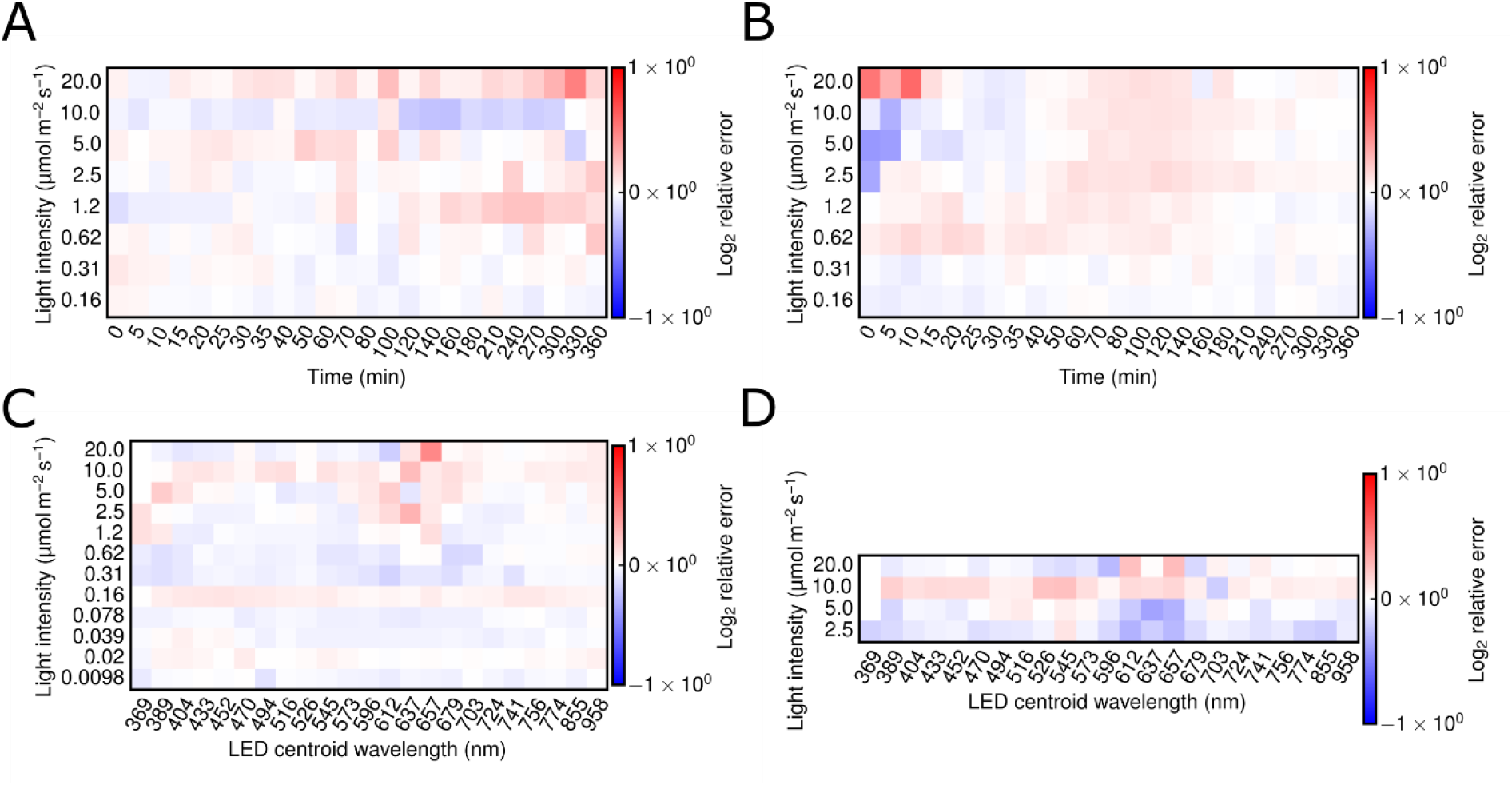
Residuals of Cph8-OmpR model to the training data. Residuals between the data and the model are shown for the (A) activating and (B) deactivating step-responses as well as the (C) forward and (D) reverse spectral measurements. The residuals are expressed relative to the measured fluorescence data on a log (base 2) scale (i.e. log_2_ *F*_data_/*F*_model_.

**Fig S12.**
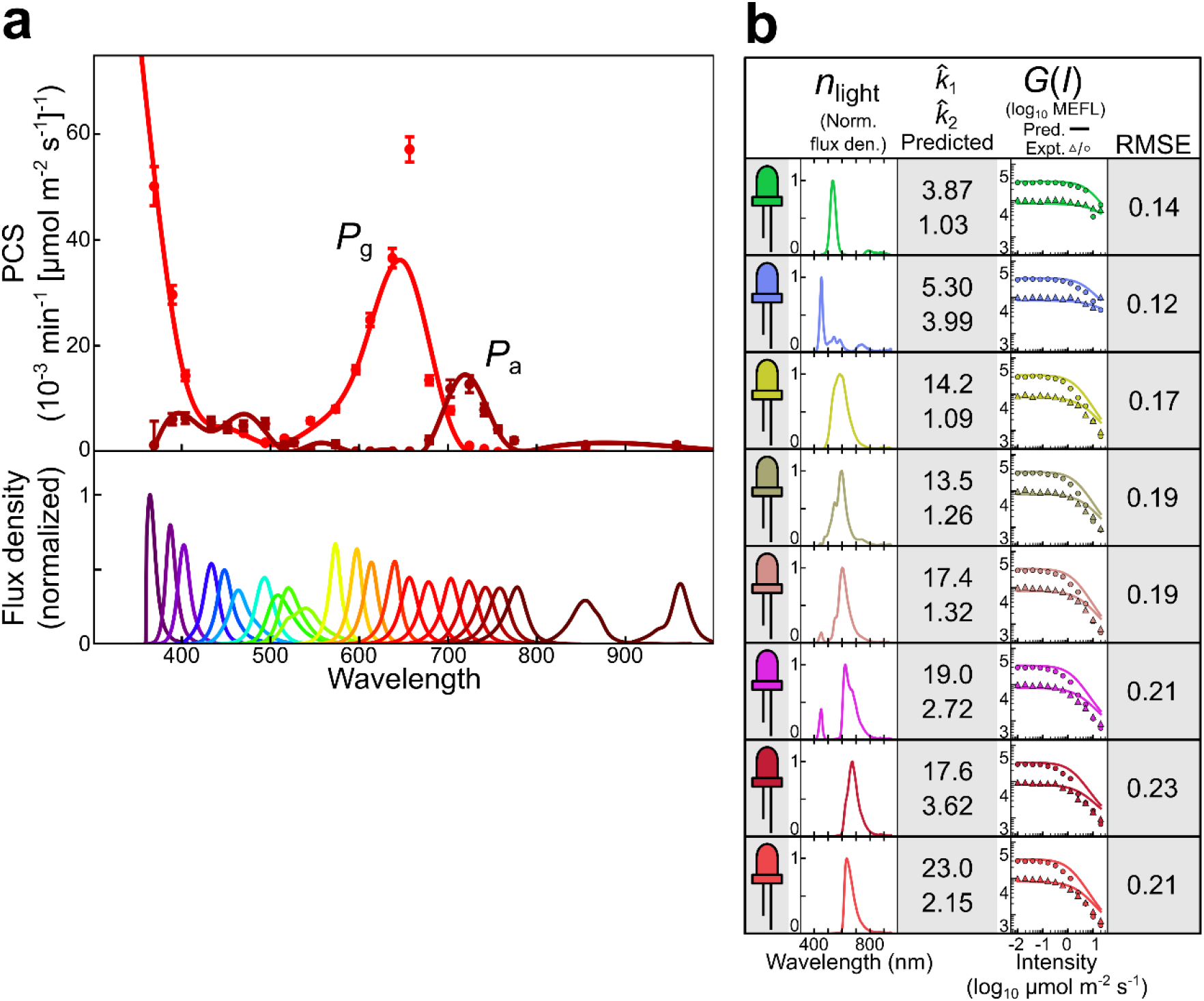
Cph8 PCS estimation and Cph8-OmpR model spectral validation. Figure details are described in Fig. 3.

**Fig S13.**
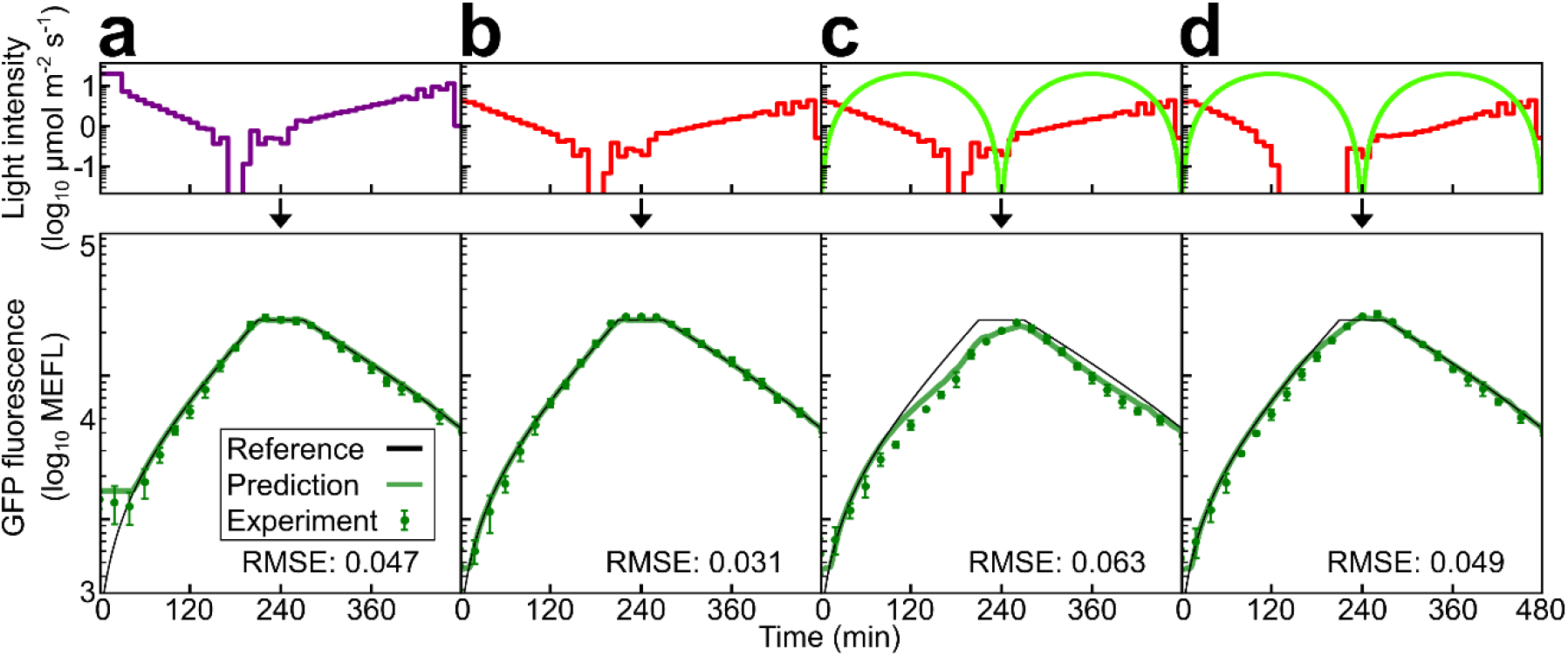
Cph8-OmpR model dynamic programming validation. Figure details are described in Fig. 4.

**Fig S14.**
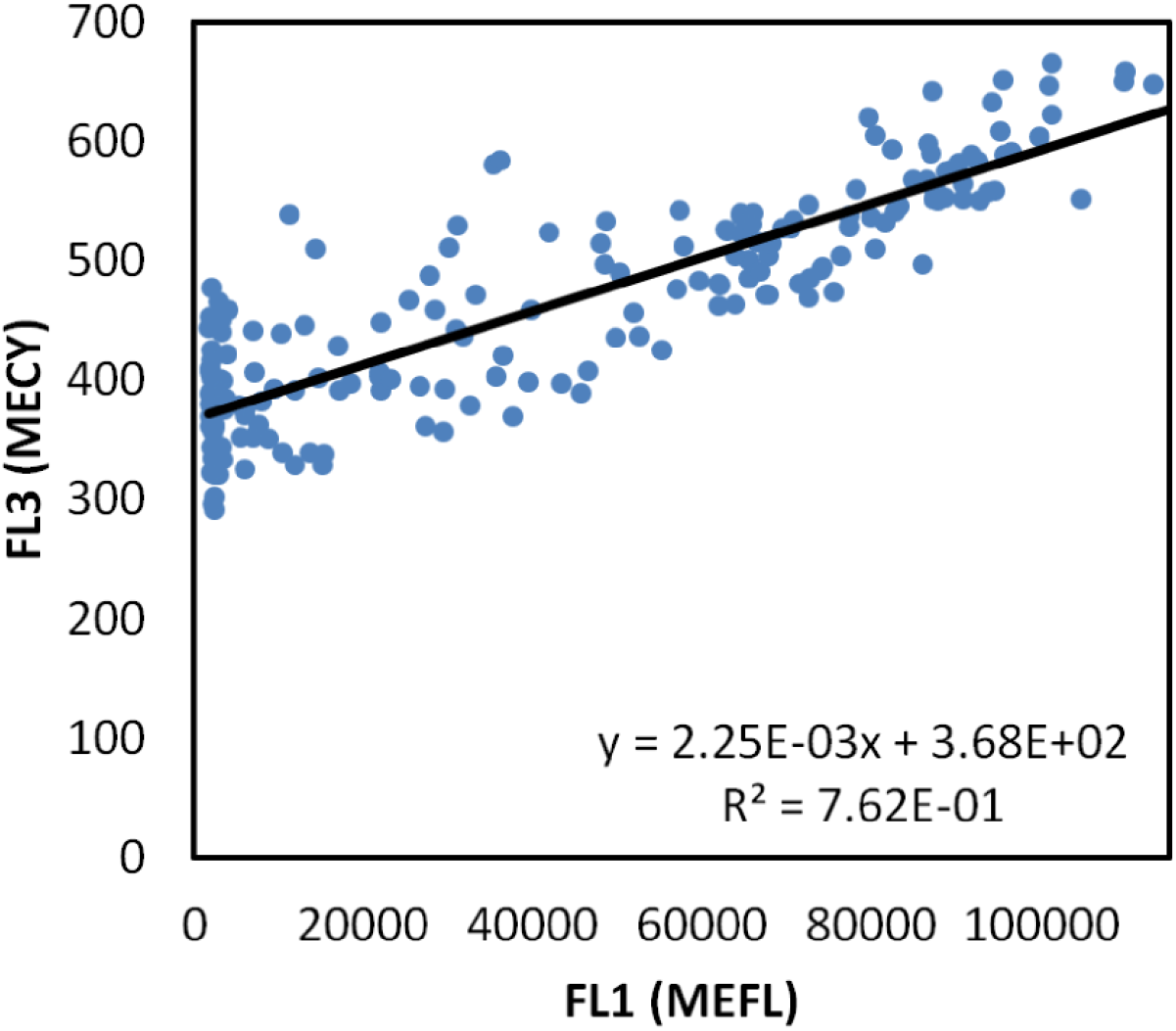
Dual-system fluorescent reporter bleedthrough compensation. High intracellular concentrations of sfGFP produce signal in the red-shifted long-pass filter cytometer channel (FL3) typically used for mCherry readout. To compensate for this bleedthrough, the “rgv2_r01” experimental data (**File S1**), containing a wide range of sfGFP expression levels (and no mCherry) was analyzed and a linear relationship between FL3 and FL1 (the green-shifted channel used for sfGFP) was identified. A linear fit to the data was performed, and the fit is used to compensate and blank the FL3 channel to enable quantification of the mCherry signal. The compensation function used was *mCherry =* FL3 — 368 MECY- FL1 · 2.25 × 10^−3^MECY/MEFL.

**Fig S15.**
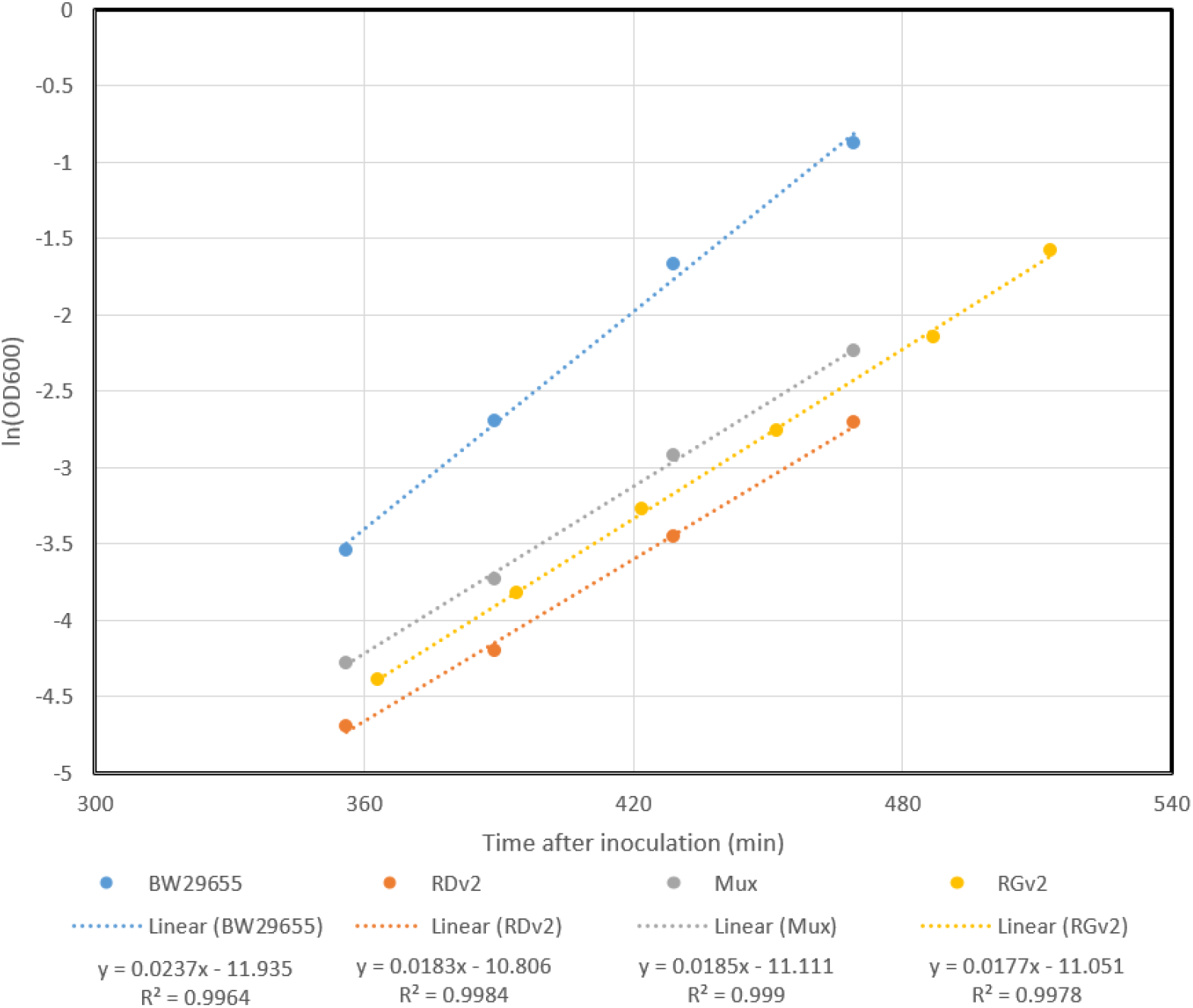
Optogenetic strain growth rate measurements. Growth rates of cells containing no plasmids (BW29655), the Cph8-OmpR system (RDv2), the dual-system (Mux), and the CcaSR system (RGv2) were measured by inoculating 8 mL M9 cultures (with appropriate antibiotics, i.e. experimental media) to the initial densities used for experiments (**Note S1**) and later drawing 1 mL samples from these cultures for OD600 measurements in a Cary 50 spectrophotometer. Linear fits were made to semi-logged data in order to extract the exponential growth rates of the strains (linear fits and R^2^ values shown below graph).

**Fig S16.**
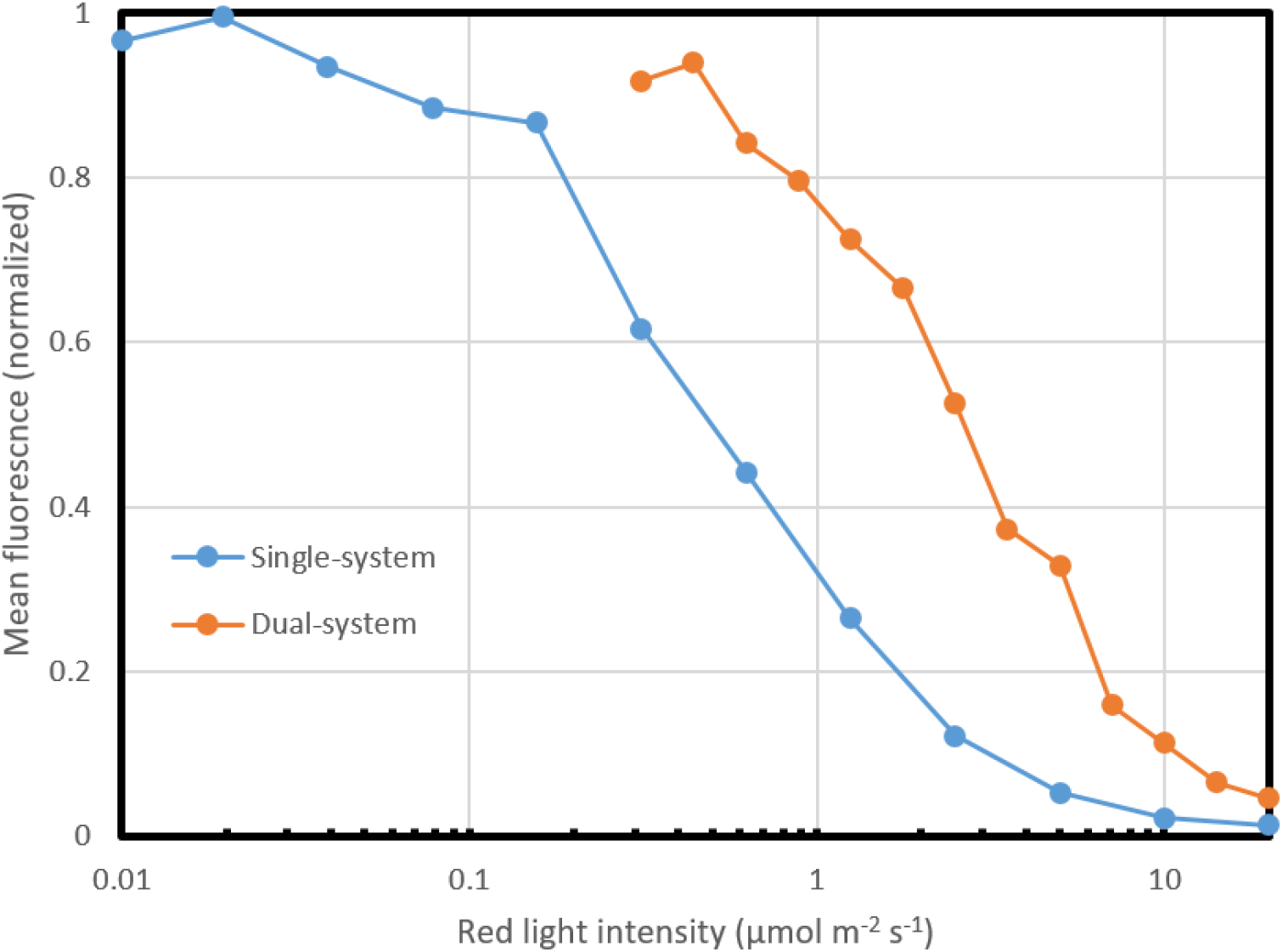
Comparison of response of single- vs. dual-system Cph8-OmpR to red light. The decreased sensitivity of the dual-system to red light is made apparent when the normalized responses of the single and dual-system Cph8-OmpR are compared. The normalization is made using the best-fit *a* and *b* values for each system in order to account for the different units and different reporters used by these systems (sfGFP for the single-strain and mCherry for the dualstrain).

**Table S1.** Manufacturer information for LEDs and filters used in this study. All data shown is pulled from manufacturer data sheets. “LED ID” is the custom identifier we have given to each LED, “SBL” is “Super Bright LEDs”, “V_drop” is the forward voltage drop across the LED, “I_test” is the test current used by the manufacturer while making reference measurements, “Inten.” is the intensity reported in units of “Inten. Units”, and “Filter ID” corresponds to the filter identification number provided by the filter manufacturer (Rosco).

| LED ID | Manufacturer | Mfg. Model | V_drop (V) | I_test (mA) | Centroid (nm) | Peak (nm) | Inten. | Inten. Units | Filter ID |
| --- | --- | --- | --- | --- | --- | --- | --- | --- | --- |
| 361-LS | LEDsupply | L5-0-U5TH15-1 | 3.8 | 20 | ND | 361 | 750 | uW | None |
| 380-SB | SBL | RL5-UV0230-380 | 3.5 | 20 | 380 | ND | 40 | mW | None |
| 405-SB | SBL | RL5-UV0430-400 | 3.5 | 20 | 400 | ND | 40 | mW | None |
| 430-MB | Marubeni | L430-03 | 3.4 | 20 | ND | 430 | 21 | mW | None |
| 450-MB | Marubeni | L450-03 | 3.4 | 20 | ND | 450 | 20 | mW | None |
| 470-SB | SBL | RL5-B2545 | 3.5 | 20 | 470 | 472 | 2500 | mcd | None |
| 490-MB_R383 | Marubeni | L490-03 | 3.3 | 20 | 496 | 490 | 12 | mW | R383 |
| 505-SB | SBL | RL5-A9018 | 3.6 | 20 | 505 | ND | 9000 | mcd | None |
| 520-2-KB | Kingbright | WP7083ZGD/G | 4 | 20 | 525 | 520 | 2200 | mcd | None |
| 525-SB_R2003 | SBL | RL5-G8045 | 3.5 | 20 | ND | 525 | 1600 | mcd | R2003 |
| 570-KB | Kingbright | WP7113CGCK | 2.5 | 20 | 570 | 574 | 700 | mcd | None |
| 590-SB | SBL | RL5-Y3545 | 2.4 | 20 | ND | 588 | 3500 | mcd | None |
| 605-SB | SBL | RL5-O4030 | 2 | 20 | ND | 605 | 4000 | mcd | None |
| 630-SB | SBL | RL5-R3545 | 2.2 | 20 | ND | 628 | 3500 | mcd | None |
| 660-LS | LEDsupply | L2-0-R5TH50-1 | 2.2 | 20 | ND | 660 | 2000 | mcd | None |
| 680-MT | Marktech | MTE6800N2-UR | 1.8 | 20 | ND | 680 | 5.5 | mW | None |
| 700-MB | Marubeni | L700-03AU | 2 | 50 | ND | 700 | 14 | mW | None |
| 720-MB | Marubeni | L720-03-AU | 1.8 | 50 | ND | 720 | 13 | mW | None |
| 740-MT | Marktech | MTE1074N1-R | 1.8 | 20 | ND | 740 | 4 | mW | None |
| 760-MB | Marubeni | L760-04-AU | 1.8 | 50 | ND | 760 | 19 | mW | None |
| 780-MB | Marubeni | L780-04-AU | 1.6 | 50 | ND | 780 | 28 | mW | None |
| 850-VI | Vishay Infrared | TSHG6200 | 1.5 | 100 | ND | 850 | 50 | mW | None |
| 940-VI | Vishay Infrared | TSAL6400 | 1.35 | 100 | ND | 940 | 40 | mW | None |
| white | SBL | RL5-W18030 | 3.4 | 20 | ND | ND | 18000 | mcd | None |
| white_R12 | Same as above LED |  |  |  |  |  |  |  | R12 |
| white_R27 |  |  |  |  |  |  |  |  | R27 |
| white_R39 |  |  |  |  |  |  |  |  | R39 |
| white_R90 |  |  |  |  |  |  |  |  | R90 |
| white_R120 |  |  |  |  |  |  |  |  | R120 |
| white_R2007 |  |  |  |  |  |  |  |  | R2007 |
| white_R3150 |  |  |  |  |  |  |  |  | R3150 |
| white_3310 |  |  |  |  |  |  |  |  | R3310 |

**Table S2.**
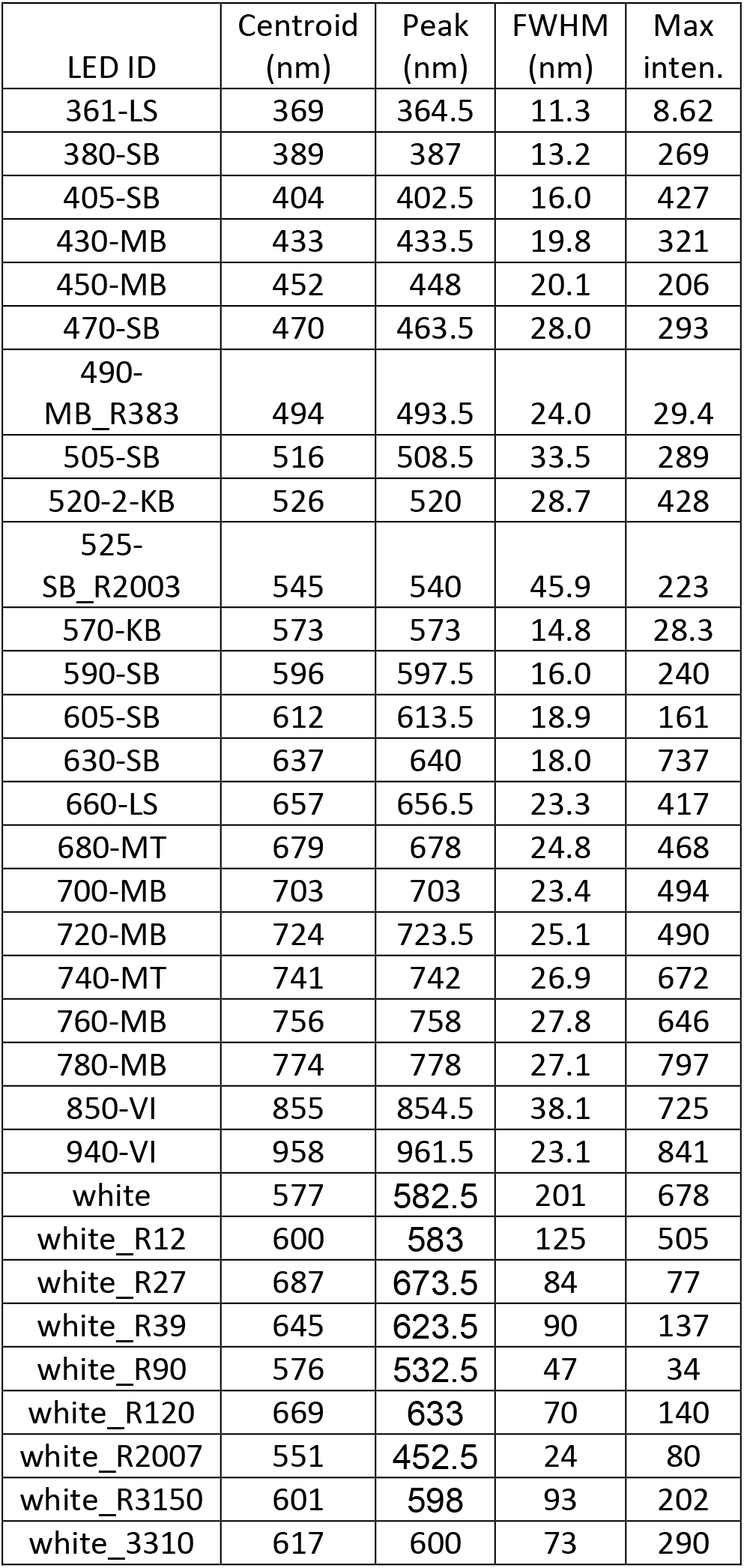
LED reference measurements. Reference measurements were made for each LED by powering the LED with the maximum LPA current (20 mA) and the minimum spectroradiometer integration time necessary to achieve 80% saturation of the detector in order to maximize the signal-to-noise ratio of the measurement. Statistics generated from these reference spectral measurements are shown in this table (“FWHM” is “Full-Width at Half-Maximum”, “Max inten.” is in units of µmol m^−2^ s^−1^.

**Table S3.**
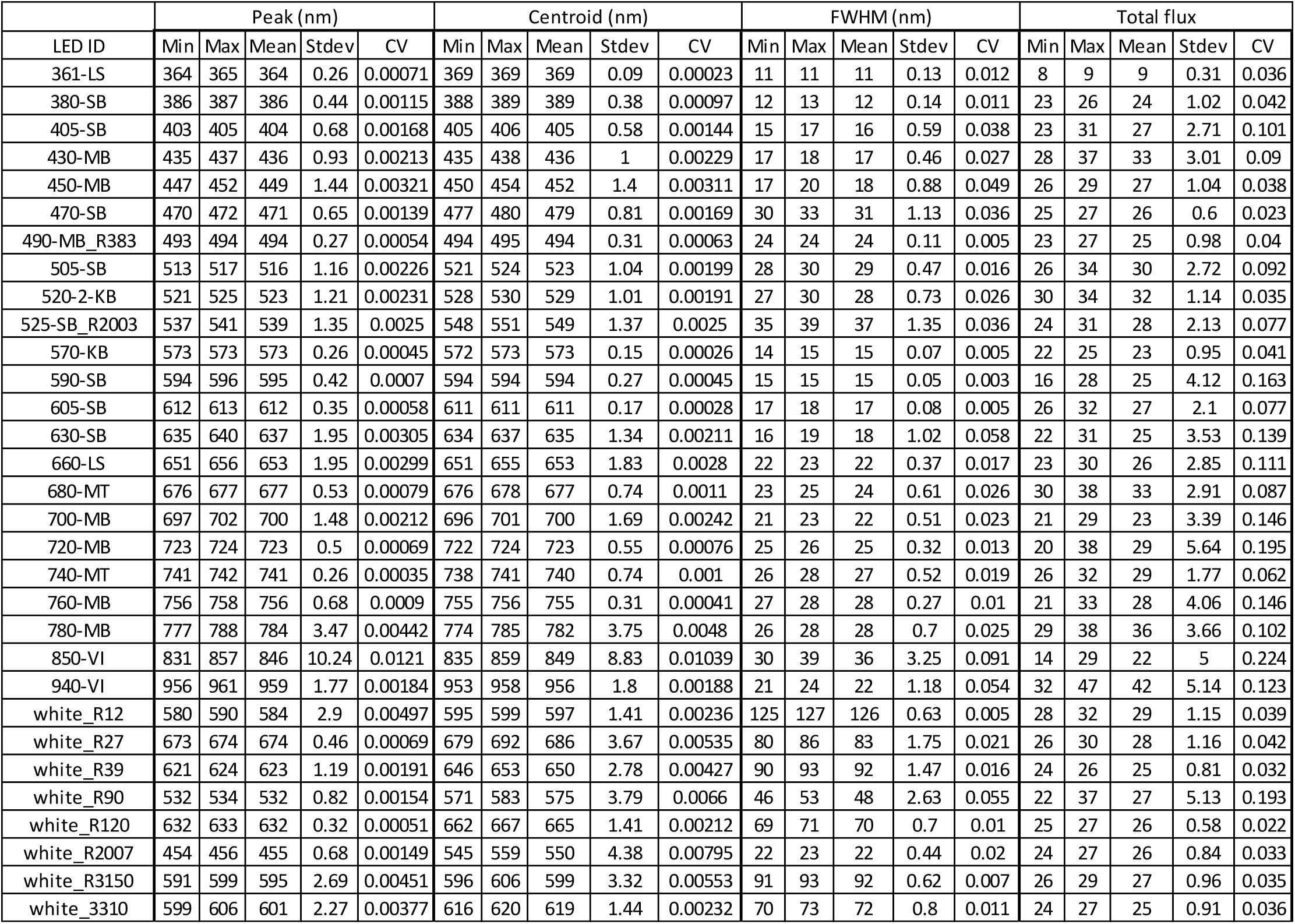
LED calibration results. Statistics generated during the calibration of the LEDs used in this study are shown. These results were calculated after the first-pass of calibration and thus are representative of the variation present in each set of LEDs given the same settings in the LPA (i.e. all LEDs used in each row of the table had the same “dot-correction” and “gray-scale calibration” value).

**Table S4.** Constraints and initial values for CcaSR model regression.

|  |  | Round 1 |  | Round 2 |  | Result |  |
| --- | --- | --- | --- | --- | --- | --- | --- |
| Parameter | Unit | Init. value | Bounds | Init. value | Bounds | Value | Std. err. |
| $b$ | MEFL | $1.5 \times 10^3$ | $[1 \times 10^{-6}, \infty]$ | $1.5 \times 10^3$ | $[1 \times 10^{-6}, \infty]$ | $1.65 \times 10^3$ | $1.50 \times 10^1$ |
| $a$ | MEFL | $1.1 \times 10^5$ | $[1 \times 10^{-6}, \infty]$ | $1.1 \times 10^5$ | $[1 \times 10^{-6}, \infty]$ | $9.74 \times 10^4$ | $1.19 \times 10^3$ |
| $n$ | | 2 | $[1 \times 10^{-6}, \infty]$ | 2 | $[1 \times 10^{-6}, \infty]$ | 2.11 | $2.55 \times 10^{-2}$ |
| $k$ | | 1 | $[1 \times 10^{-6}, \infty]$ | 0.3190063 | Fixed | $3.19 \times 10^{-1}$ | 0 |
| $k_g$ | $\text{min}^{-1}$ | $1.8515 \times 10^{-2}$ | Fixed | $1.8515 \times 10^{-2}$ | Fixed | $1.85 \times 10^{-2}$ | 0 |
| $k_{dr}$ | $\text{min}^{-1}$ | 0.1 | $[1 \times 10^{-6}, \infty]$ | 0 | Fixed | 0.00 | 0 |
| $\tau$ | min | 10 | [5, 20] | 10 | [5, 20] | $1.15 \times 10^1$ | $2.71 \times 10^{-1}$ |
| Parameter | Unit | Init. value | Bounds | Init. value | Bounds | Value | Std. err. |
| $\hat{k}_{1,0}$ | $\text{min}^{-1} / (\mu\text{mol}/\text{m}^2/\text{s})$ | $1 \times 10^{-2}$ | $[1 \times 10^{-6}, \infty]$ | $1 \times 10^{-2}$ | $[1 \times 10^{-6}, \infty]$ | $1.34 \times 10^{-2}$ | $9.44 \times 10^{-4}$ |
| $\hat{k}_{1,1}$ | $\text{min}^{-1} / (\mu\text{mol}/\text{m}^2/\text{s})$ | $1 \times 10^{-2}$ | $[1 \times 10^{-6}, \infty]$ | $1 \times 10^{-2}$ | $[1 \times 10^{-6}, \infty]$ | $8.40 \times 10^{-3}$ | $5.92 \times 10^{-4}$ |
| $\hat{k}_{1,2}$ | $\text{min}^{-1} / (\mu\text{mol}/\text{m}^2/\text{s})$ | $1 \times 10^{-2}$ | $[1 \times 10^{-6}, \infty]$ | $1 \times 10^{-2}$ | $[1 \times 10^{-6}, \infty]$ | $4.80 \times 10^{-3}$ | $3.24 \times 10^{-4}$ |
| $\hat{k}_{1,3}$ | $\text{min}^{-1} / (\mu\text{mol}/\text{m}^2/\text{s})$ | $1 \times 10^{-2}$ | $[1 \times 10^{-6}, \infty]$ | $1 \times 10^{-2}$ | $[1 \times 10^{-6}, \infty]$ | $3.09 \times 10^{-3}$ | $2.16 \times 10^{-4}$ |
| $\hat{k}_{1,4}$ | $\text{min}^{-1} / (\mu\text{mol}/\text{m}^2/\text{s})$ | $1 \times 10^{-2}$ | $[1 \times 10^{-6}, \infty]$ | $1 \times 10^{-2}$ | $[1 \times 10^{-6}, \infty]$ | $5.90 \times 10^{-3}$ | $4.14 \times 10^{-4}$ |
| $\hat{k}_{1,5}$ | $\text{min}^{-1} / (\mu\text{mol}/\text{m}^2/\text{s})$ | $1 \times 10^{-2}$ | $[1 \times 10^{-6}, \infty]$ | $1 \times 10^{-2}$ | $[1 \times 10^{-6}, \infty]$ | $1.19 \times 10^{-2}$ | $5.83 \times 10^{-4}$ |
| $\hat{k}_{1,6}$ | $\text{min}^{-1} / (\mu\text{mol}/\text{m}^2/\text{s})$ | $1 \times 10^{-2}$ | $[1 \times 10^{-6}, \infty]$ | $1 \times 10^{-2}$ | $[1 \times 10^{-6}, \infty]$ | $9.85 \times 10^{-3}$ | $4.99 \times 10^{-4}$ |
| $\hat{k}_{1,7}$ | $\text{min}^{-1} / (\mu\text{mol}/\text{m}^2/\text{s})$ | $1 \times 10^{-2}$ | $[1 \times 10^{-6}, \infty]$ | $1 \times 10^{-2}$ | $[1 \times 10^{-6}, \infty]$ | $1.07 \times 10^{-2}$ | $5.49 \times 10^{-4}$ |
| $\hat{k}_{1,8}$ | $\text{min}^{-1} / (\mu\text{mol}/\text{m}^2/\text{s})$ | $1 \times 10^{-2}$ | $[1 \times 10^{-6}, \infty]$ | $1 \times 10^{-2}$ | $[1 \times 10^{-6}, \infty]$ | $6.68 \times 10^{-3}$ | $1.13 \times 10^{-4}$ |
| $\hat{k}_{1,9}$ | $\text{min}^{-1} / (\mu\text{mol}/\text{m}^2/\text{s})$ | $1 \times 10^{-2}$ | $[1 \times 10^{-6}, \infty]$ | $1 \times 10^{-2}$ | $[1 \times 10^{-6}, \infty]$ | $1.15 \times 10^{-2}$ | $5.74 \times 10^{-4}$ |
| $\hat{k}_{1,10}$ | $\text{min}^{-1} / (\mu\text{mol}/\text{m}^2/\text{s})$ | $1 \times 10^{-2}$ | $[1 \times 10^{-6}, \infty]$ | $1 \times 10^{-2}$ | $[1 \times 10^{-6}, \infty]$ | $6.30 \times 10^{-3}$ | $4.44 \times 10^{-4}$ |
| $\hat{k}_{1,11}$ | $\text{min}^{-1} / (\mu\text{mol}/\text{m}^2/\text{s})$ | $1 \times 10^{-2}$ | $[1 \times 10^{-6}, \infty]$ | $1 \times 10^{-2}$ | $[1 \times 10^{-6}, \infty]$ | $1.92 \times 10^{-3}$ | $1.39 \times 10^{-4}$ |
| $\hat{k}_{1,12}$ | $\text{min}^{-1} / (\mu\text{mol}/\text{m}^2/\text{s})$ | $1 \times 10^{-2}$ | $[1 \times 10^{-6}, \infty]$ | $1 \times 10^{-2}$ | $[1 \times 10^{-6}, \infty]$ | $5.25 \times 10^{-4}$ | $5.09 \times 10^{-5}$ |
| $\hat{k}_{1,13}$ | $\text{min}^{-1} / (\mu\text{mol}/\text{m}^2/\text{s})$ | $1 \times 10^{-3}$ | $[1 \times 10^{-6}, \infty]$ | $1 \times 10^{-6}$ | Fixed | $1.00 \times 10^{-6}$ | 0 |
| $\hat{k}_{1,14}$ | $\text{min}^{-1} / (\mu\text{mol}/\text{m}^2/\text{s})$ | $1 \times 10^{-3}$ | $[1 \times 10^{-6}, \infty]$ | $1 \times 10^{-6}$ | Fixed | $1.00 \times 10^{-6}$ | 0 |
| $\hat{k}_{1,15}$ | $\text{min}^{-1} / (\mu\text{mol}/\text{m}^2/\text{s})$ | $1 \times 10^{-3}$ | $[1 \times 10^{-6}, \infty]$ | $1 \times 10^{-6}$ | Fixed | $1.00 \times 10^{-6}$ | 0 |
| $\hat{k}_{1,16}$ | $\text{min}^{-1} / (\mu\text{mol}/\text{m}^2/\text{s})$ | $1 \times 10^{-3}$ | $[1 \times 10^{-6}, \infty]$ | $1 \times 10^{-6}$ | Fixed | $1.00 \times 10^{-6}$ | 0 |
| $\hat{k}_{1,17}$ | $\text{min}^{-1} / (\mu\text{mol}/\text{m}^2/\text{s})$ | $1 \times 10^{-3}$ | $[1 \times 10^{-6}, \infty]$ | $1 \times 10^{-6}$ | Fixed | $1.00 \times 10^{-6}$ | 0 |
| $\hat{k}_{1,18}$ | $\text{min}^{-1} / (\mu\text{mol}/\text{m}^2/\text{s})$ | $1 \times 10^{-3}$ | $[1 \times 10^{-6}, \infty]$ | $1 \times 10^{-6}$ | Fixed | $1.00 \times 10^{-6}$ | 0 |
| $\hat{k}_{1,19}$ | $\text{min}^{-1} / (\mu\text{mol}/\text{m}^2/\text{s})$ | $1 \times 10^{-3}$ | $[1 \times 10^{-6}, \infty]$ | $1 \times 10^{-6}$ | Fixed | $1.00 \times 10^{-6}$ | 0 |
| $\hat{k}_{1,20}$ | $\text{min}^{-1} / (\mu\text{mol}/\text{m}^2/\text{s})$ | $1 \times 10^{-3}$ | $[1 \times 10^{-6}, \infty]$ | $1 \times 10^{-6}$ | Fixed | $1.00 \times 10^{-6}$ | 0 |
| $\hat{k}_{1,21}$ | $\text{min}^{-1} / (\mu\text{mol}/\text{m}^2/\text{s})$ | $1 \times 10^{-3}$ | $[1 \times 10^{-6}, \infty]$ | $1 \times 10^{-6}$ | Fixed | $1.00 \times 10^{-6}$ | 0 |
| $\hat{k}_{1,22}$ | $\text{min}^{-1} / (\mu\text{mol}/\text{m}^2/\text{s})$ | $1 \times 10^{-3}$ | $[1 \times 10^{-6}, \infty]$ | $1 \times 10^{-6}$ | Fixed | $1.00 \times 10^{-6}$ | 0 |

**Table S4.** Constraints and initial values for CcaSR model regression.
| Parameter | Unit | Round 1 |  | Round 2 |  | Result |  |
| --- | --- | --- | --- | --- | --- | --- | --- |
|  |  | Init. value | Bounds | Init. value | Bounds | Value | Std. err. |
| $\hat{k}_{2,0}$ | $\text{min}^{-1}/(\mu\text{mol}/\text{m}^2/\text{s})$ | $1 \times 10^{-3}$ | $[1 \times 10^{-6}, \infty]$ | $1 \times 10^{-3}$ | $[1 \times 10^{-6}, \infty]$ | $1.26 \times 10^{-2}$ | $4.09 \times 10^{-3}$ |
| $\hat{k}_{2,1}$ | $\text{min}^{-1}/(\mu\text{mol}/\text{m}^2/\text{s})$ | $1 \times 10^{-3}$ | $[1 \times 10^{-6}, \infty]$ | $1 \times 10^{-3}$ | $[1 \times 10^{-6}, \infty]$ | $1.69 \times 10^{-3}$ | $2.68 \times 10^{-3}$ |
| $\hat{k}_{2,2}$ | $\text{min}^{-1}/(\mu\text{mol}/\text{m}^2/\text{s})$ | $1 \times 10^{-3}$ | $[1 \times 10^{-6}, \infty]$ | $1 \times 10^{-3}$ | $[1 \times 10^{-6}, \infty]$ | $5.44 \times 10^{-3}$ | $1.22 \times 10^{-3}$ |
| $\hat{k}_{2,3}$ | $\text{min}^{-1}/(\mu\text{mol}/\text{m}^2/\text{s})$ | $1 \times 10^{-3}$ | $[1 \times 10^{-6}, \infty]$ | $1 \times 10^{-3}$ | $[1 \times 10^{-6}, \infty]$ | $1.51 \times 10^{-3}$ | $9.47 \times 10^{-4}$ |
| $\hat{k}_{2,4}$ | $\text{min}^{-1}/(\mu\text{mol}/\text{m}^2/\text{s})$ | $1 \times 10^{-3}$ | $[1 \times 10^{-6}, \infty]$ | $1 \times 10^{-3}$ | $[1 \times 10^{-6}, \infty]$ | $1.89 \times 10^{-3}$ | $1.83 \times 10^{-3}$ |
| $\hat{k}_{2,5}$ | $\text{min}^{-1}/(\mu\text{mol}/\text{m}^2/\text{s})$ | $1 \times 10^{-3}$ | $[1 \times 10^{-6}, \infty]$ | $1 \times 10^{-6}$ | Fixed | $1.00 \times 10^{-6}$ | 0 |
| $\hat{k}_{2,6}$ | $\text{min}^{-1}/(\mu\text{mol}/\text{m}^2/\text{s})$ | $1 \times 10^{-3}$ | $[1 \times 10^{-6}, \infty]$ | $1 \times 10^{-6}$ | Fixed | $1.00 \times 10^{-6}$ | 0 |
| $\hat{k}_{2,7}$ | $\text{min}^{-1}/(\mu\text{mol}/\text{m}^2/\text{s})$ | $1 \times 10^{-3}$ | $[1 \times 10^{-6}, \infty]$ | $1 \times 10^{-6}$ | Fixed | $1.00 \times 10^{-6}$ | 0 |
| $\hat{k}_{2,8}$ | $\text{min}^{-1}/(\mu\text{mol}/\text{m}^2/\text{s})$ | $1 \times 10^{-3}$ | $[1 \times 10^{-6}, \infty]$ | $1 \times 10^{-3}$ | $[1 \times 10^{-6}, \infty]$ | $8.66 \times 10^{-4}$ | $3.02 \times 10^{-4}$ |
| $\hat{k}_{2,9}$ | $\text{min}^{-1}/(\mu\text{mol}/\text{m}^2/\text{s})$ | $1 \times 10^{-3}$ | $[1 \times 10^{-6}, \infty]$ | $1 \times 10^{-6}$ | Fixed | $1.00 \times 10^{-6}$ | 0 |
| $\hat{k}_{2,10}$ | $\text{min}^{-1}/(\mu\text{mol}/\text{m}^2/\text{s})$ | $1 \times 10^{-3}$ | $[1 \times 10^{-6}, \infty]$ | $1 \times 10^{-3}$ | $[1 \times 10^{-6}, \infty]$ | $1.78 \times 10^{-3}$ | $1.94 \times 10^{-3}$ |
| $\hat{k}_{2,11}$ | $\text{min}^{-1}/(\mu\text{mol}/\text{m}^2/\text{s})$ | $1 \times 10^{-3}$ | $[1 \times 10^{-6}, \infty]$ | $1 \times 10^{-3}$ | $[1 \times 10^{-6}, \infty]$ | $2.69 \times 10^{-3}$ | $5.85 \times 10^{-4}$ |
| $\hat{k}_{2,12}$ | $\text{min}^{-1}/(\mu\text{mol}/\text{m}^2/\text{s})$ | $1 \times 10^{-3}$ | $[1 \times 10^{-6}, \infty]$ | $1 \times 10^{-3}$ | $[1 \times 10^{-6}, \infty]$ | $3.36 \times 10^{-3}$ | $4.39 \times 10^{-4}$ |
| $\hat{k}_{2,13}$ | $\text{min}^{-1}/(\mu\text{mol}/\text{m}^2/\text{s})$ | $1 \times 10^{-2}$ | $[1 \times 10^{-6}, \infty]$ | $1 \times 10^{-2}$ | $[1 \times 10^{-6}, \infty]$ | $2.98 \times 10^{-3}$ | $2.35 \times 10^{-4}$ |
| $\hat{k}_{2,14}$ | $\text{min}^{-1}/(\mu\text{mol}/\text{m}^2/\text{s})$ | $1 \times 10^{-2}$ | $[1 \times 10^{-6}, \infty]$ | $1 \times 10^{-2}$ | $[1 \times 10^{-6}, \infty]$ | $4.58 \times 10^{-3}$ | $3.11 \times 10^{-4}$ |
| $\hat{k}_{2,15}$ | $\text{min}^{-1}/(\mu\text{mol}/\text{m}^2/\text{s})$ | $1 \times 10^{-2}$ | $[1 \times 10^{-6}, \infty]$ | $1 \times 10^{-2}$ | $[1 \times 10^{-6}, \infty]$ | $1.62 \times 10^{-3}$ | $1.75 \times 10^{-4}$ |
| $\hat{k}_{2,16}$ | $\text{min}^{-1}/(\mu\text{mol}/\text{m}^2/\text{s})$ | $1 \times 10^{-2}$ | $[1 \times 10^{-6}, \infty]$ | $1 \times 10^{-2}$ | $[1 \times 10^{-6}, \infty]$ | $2.18 \times 10^{-3}$ | $1.90 \times 10^{-4}$ |
| $\hat{k}_{2,17}$ | $\text{min}^{-1}/(\mu\text{mol}/\text{m}^2/\text{s})$ | $1 \times 10^{-2}$ | $[1 \times 10^{-6}, \infty]$ | $1 \times 10^{-2}$ | $[1 \times 10^{-6}, \infty]$ | $5.62 \times 10^{-4}$ | $1.40 \times 10^{-4}$ |
| $\hat{k}_{2,18}$ | $\text{min}^{-1}/(\mu\text{mol}/\text{m}^2/\text{s})$ | $1 \times 10^{-2}$ | $[1 \times 10^{-6}, \infty]$ | $1 \times 10^{-2}$ | $[1 \times 10^{-6}, \infty]$ | $2.56 \times 10^{-4}$ | $1.50 \times 10^{-4}$ |
| $\hat{k}_{2,19}$ | $\text{min}^{-1}/(\mu\text{mol}/\text{m}^2/\text{s})$ | $1 \times 10^{-2}$ | $[1 \times 10^{-6}, \infty]$ | $1 \times 10^{-2}$ | $[1 \times 10^{-6}, \infty]$ | $1.00 \times 10^{-6}$ | 0.00 |
| $\hat{k}_{2,20}$ | $\text{min}^{-1}/(\mu\text{mol}/\text{m}^2/\text{s})$ | $1 \times 10^{-2}$ | $[1 \times 10^{-6}, \infty]$ | $1 \times 10^{-6}$ | Fixed | $1.00 \times 10^{-6}$ | 0 |
| $\hat{k}_{2,21}$ | $\text{min}^{-1}/(\mu\text{mol}/\text{m}^2/\text{s})$ | $1 \times 10^{-2}$ | $[1 \times 10^{-6}, \infty]$ | $1 \times 10^{-6}$ | Fixed | $1.00 \times 10^{-6}$ | 0 |
| $\hat{k}_{2,22}$ | $\text{min}^{-1}/(\mu\text{mol}/\text{m}^2/\text{s})$ | $1 \times 10^{-2}$ | $[1 \times 10^{-6}, \infty]$ | $1 \times 10^{-6}$ | Fixed | $1.00 \times 10^{-6}$ | 0 |

**Table S5.**
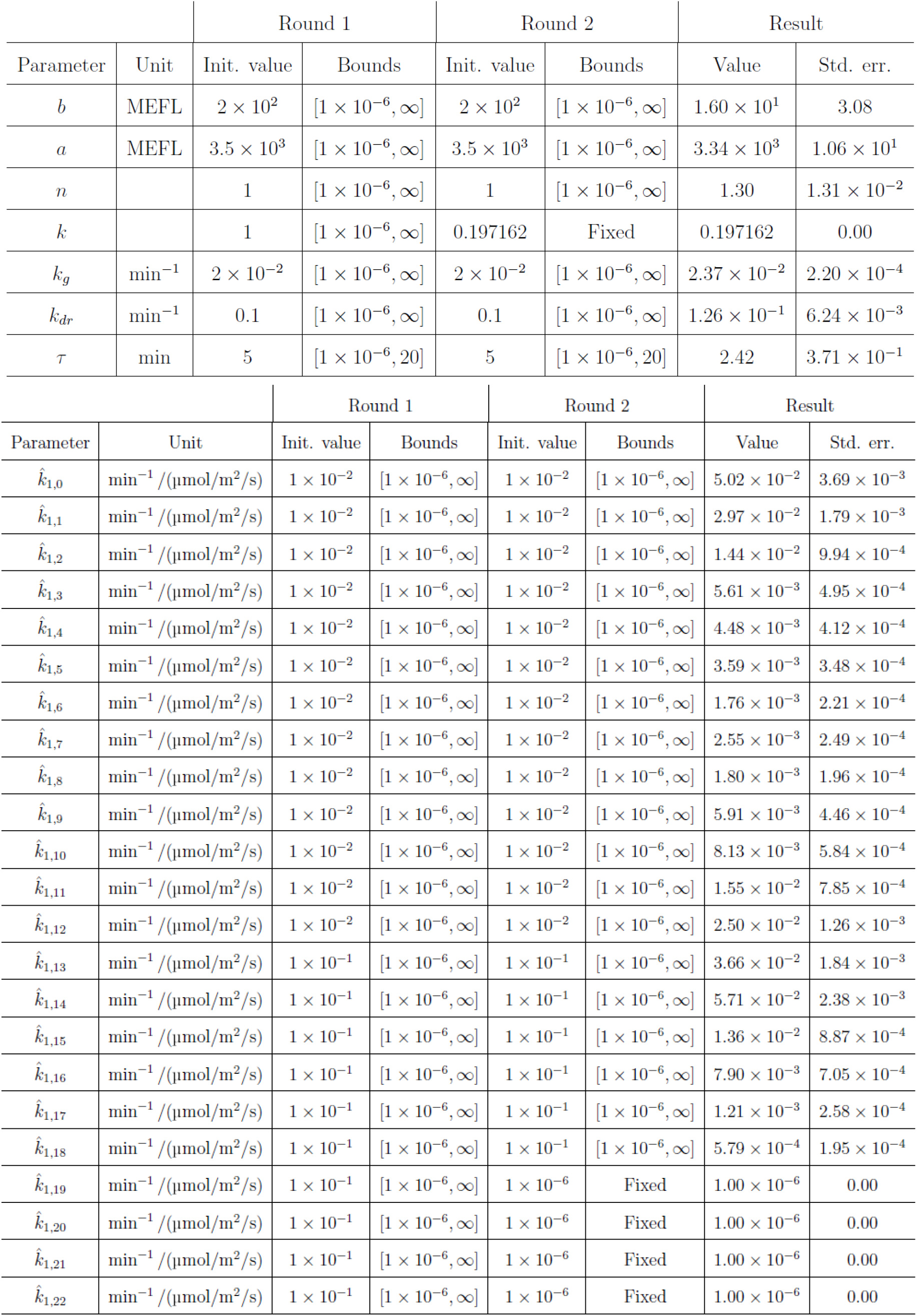
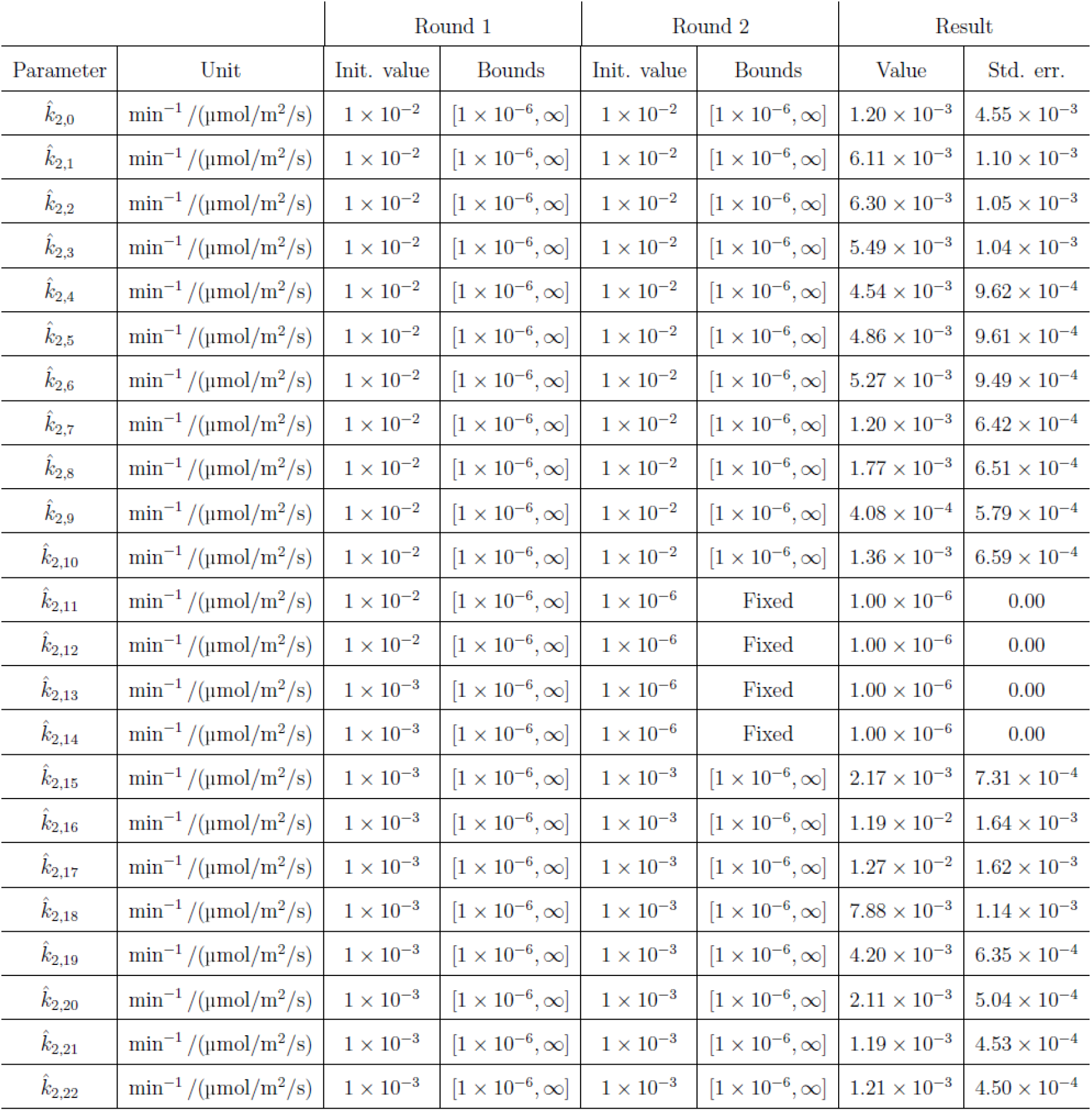
Constraints and initial values for Cph8-OmpR model regression.

**Table S6.**
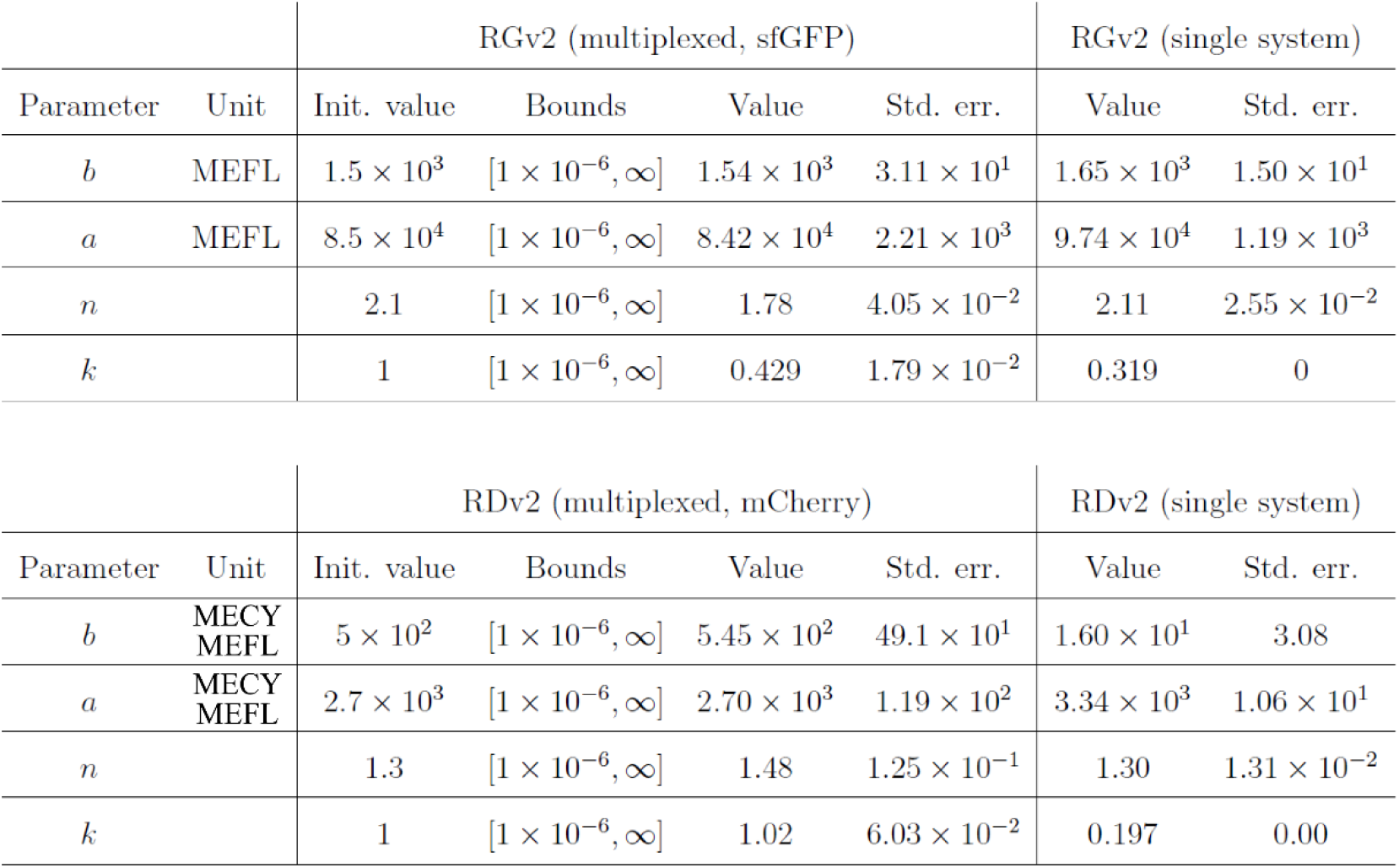
Dual-system model parameters.

**SI Files**

**File S1** Plate-by-plate overview of CcaSR experiments. This Excel spreadsheet contains a description of the light programs used for each 24-well plate used for the CcaSR experiments. On the “experiment” sheet, the “Run ID” and “Run name” columns contain unique identifiers for each experimental trial. The “Strain ID” is an identifier for the CcaSR strain. “LPA” is the name of the LPA device used. “Top LEDs” and “Bottom LEDs” correspond to the LED set identifiers used in the LED calibration archives. “Randomize” indicates whether the light program time points were randomized throughout the wells of the plate. “Program type” is an identifier corresponding to the type of light program (“aas” is a forward action spectrum, “ras” is a reverse action spectrum, “dta” is an activating step change, “atd” is a deactivating step change, “dv” is a dynamic validation, and “sv” is a spectral validation). The parameter columns further specify the details of the light program (intensities are in µmol m^−2^ s^−1^, “Model” refers to an identifier for the system model used to generate the light program, reference signals and perturbation signals are identifiers corresponding to reference programs listed in **File S3**). For dynamic validation experiments, the model, reference signal, and perturbation signal (if present) are used to generate light programs which are available in **File S4**.

**File S2** Experimental measurements. This folder contains spreadsheets detailing all experimental measurements used in this manuscript. The “rgv2” folder corresponds to CcaSR, “rdv2” to Cph8-OmpR, and “mux” to the dual-system. Within these folders are the spreadsheets, identified using the “Run ID” values indicated in **File S1,7-8**. Within each of these spreadsheets are a series of tabs corresponding to the experimental measurements (including cytometry histograms, final OD600 values, and incubator temperature time-courses) and the light programs (both in intensity units and in LPA-readable 14-bit grayscale) used to generate them. Each sample can be correlated between tabs using the “Sample ID” column, which provides a unique identifier for each culture sample.

**File S3** Reference LED spectra. The raw spectroradiometric reference measurement of each LED is available as a .IRR file (human-readable in a text editor). The post-processed .xls file contains a truncated spectrum and statistics about the LED spectrum. The “reference_leds.xls” file contains the summary statistics for each of the LEDs.

**File S4** LED calibration measurements and results. The “process_led_archives.py” Python 2.7 script parses the “led_archives.xlsx” and processes the contents of the “raw” and “sd” folders within each of the LED archive folders according to the LED layouts in the “layouts” folder. The “raw” folder contains the raw calibration measurements, while the “sd” folder contains the SD card files from the LPA used during calibration. The “processed” folder contains spreadsheets with spectra and LED statistics generated for each LED. During processing, an additional file is generated alongside the “processed” folder containing a summary of the LED calibration information for each set of LEDs.

**File S5** Reference gene expression signals. The reference programs and perturbation signals used for light program generation are stored in this file. Each reference program consists of a pair of columns identified by “[ID]_x” and “[ID]_y”. The first column contains time values (in minutes) while the values of the second column depend upon whether the signal is a reference or a perturbation (as indicated by the ID). Reference signals (ID=”ref”) describe normalized gene expression levels (i.e. 0 is the minimal output of a system, and 1 is the maximum), while perturbation signals (ID=”pert”) describe perturbative light programs as a sequence of intensities (in µmol m^−2^ s^−1^).

**File S6** Generated light programs. The light programs produced using the Light Program Generator algorithm are stored in this file. “LED ID” corresponds to the LED which follows the light program, “Times” are the time points (in minutes) corresponding to step-changes in the light intensity, “Intensities” is a sequence of intensities describing the generated light program, “Pre-inten” is the preconditioning intensity used for the program, “Ref ID” is the reference signal used to generate the program, “Model ID” is the identifier of the model used to generate the program, “Compensated perturbation LED ID” is the LED ID for the perturbing LED (if present), and “Compensated perturbation ref ID” is the perturbation signal used by the perturbing LED (if present).

**File S7** Plate-by-plate overview of Cph8-OmpR experiments. File contents follow the same description used for **File S1**.

**File S8** Plate-by-plate overview of dual-system programming experiments. File contents follow the same description used for **File S1**.

